# Thermostable *in vitro* transcription-translation for enzyme screening in microdroplets

**DOI:** 10.1101/2024.02.22.580490

**Authors:** Ana L. J. L. Ribeiro, Patricia Pérez-Arnaiz, Mercedes Sánchez-Costa, Lara Pérez, Marcos Almendros, Liisa van Vliet, Fabrice Gielen, Jesmine Lim, Simon Charnock, Florian Hollfelder, J. Eduardo González-Pastor, José Berenguer, Aurelio Hidalgo

## Abstract

**Background:** *In vitro* expression involves the utilization of the transcription and translation machinery derived from the cell to produce one or more proteins of interest and has found widespread application in the optimization of gene circuits or metabolic pathways in synthetic biology but also in pharmaceutical manufacturing. Most *in vitro* expression systems available are active at moderate temperatures but to screen large libraries of natural or artificial genetic diversity for highly thermostable enzymes or enzyme variants, it is instrumental to enable protein synthesis at high temperatures. Moreover, given the fact that the main barrier toward the general use of *in vitro* expression is its high price compared with host-based recombinant expression, there is a need to develop alternative *in vitro* expression systems operating at high temperatures and compatible with technologies that enable ultrahigh-throughput screening in reduced volumes, such as microfluidic water-in-oil (w/o) droplets.

**Results:** To this end, we produced high-expression cell-free extracts from *Thermus thermophilus* for *in vitro* translation and supplemented them with thermostable enzymatic cascades for energy regeneration and a moderately thermostable RNA polymerase for transcription, which ultimately limited the temperature of protein synthesis. The yield was comparable to other thermostable *in vitro* expression systems, while the preparation procedure is simpler and can be suited to different *Thermus thermophilus* strains. Furthermore, these extracts have enabled in vitro expression in microfluidic droplets at high temperatures for the first time. Although the composition of these extracts showed a high background in carboxyl esterase assays, β-glucosidase and cellobiose hydrolase activities could be measured with minimal background.

**Conclusions:** Cell-free extracts from *Thermus thermophilus* represent a simpler alternative to heavily optimized or pure component thermostable *in vitro* expression systems. Moreover, due to their compatibility with droplet microfluidics and enzyme assays at high temperatures, the reported system represents a convenient gateway for enzyme screening at higher temperatures with ultrahigh-throughput.

## BACKGROUND

Cell-free protein synthesis (CFPS) involves the utilization of the transcription and translation machinery of the cell to produce proteins of interest, independently of constraints imposed by cellular viability and variability as well as membrane integrity (1). This acellular paradigm allows a higher tolerance for toxic substrates or products, easy manipulation and a more precise control over both the synthetic components and the synthesized products, especially in complex biological networks, such as gene circuits or metabolic pathways (2–4). In fact, *in vitro* cell-free systems allow a more precise control over the experimental parameters and the customization of one or more steps in the workflow of protein synthesis even at high-throughput, such as transcription, translation or post-translational modifications. For instance, CFPS allows a straightforward incorporation of unnatural amino acids to add (non-)biological functional groups into proteins for site-directed, click chemistry-based labelling (5), photoactivation (6), redox groups to study and engineer electron transfer processes in proteins (7) or for post-translational modifications like histone acetylation (8), histone methylation (9), ubiquitination (10) and incorporation of phosphoserine residues (11), glycans and metal centers (12,13), among others. Finally, cell-free systems have also been used for the incorporation of selenomethionine residues in proteins for X-ray crystallography structure determination (14).

Whereas *in vivo*, protein synthesis is coupled to the cellular ATP/GTP pools, in a cell-free context these energy-rich molecules must be provided by very expensive, high-energy phosphate compounds and auxiliary enzymes, such as those involved in glycolysis and the Krebs cycle. In spite of this limitation and mostly owing to its reproducibility and controlled environment, *in vitro* protein synthesis has rapidly found its way from basic research into application. Noteworthy uses of CFPS in basic research include the synthesis of protein libraries for functional genomics and structural biology, biological network prototyping by accelerating the design-build-test cycle (15), artificial cell systems (16–18), miniaturization and protein arrays (19,20), diagnostics and biosensors (21–23), prototyping of biosynthetic pathways and metabolic engineering (24–26) and the production of functional membrane proteins (27–29), whereas one of the most outstanding applications is the production of biopharmaceuticals such as antimicrobial peptides (30), cytokines (31), vaccines (32) and antibodies (33).

Several cell-free expression systems have been developed, the most well-established being from *Escherichia coli* (34), wheat germ (35,36), yeast (37,38) rabbit reticulocyte (39) or HeLa cells (40), but further examples include extracts derived from *Streptomyces venezuelae* (41,42), *Bacillus subtilis* (43), *Vibrio natriegens* (44–46), insect *Spodoptera frugiperda* 21 (*Sf*21) (47,48), Chinese hamster ovary (49), tobacco BY-2 (50) and *Leishmania tarentolae* (51,52). Each of the mentioned CFPS systems has different advantages and limitations and therefore, the choice of source depends on the nature of the protein to be expressed and downstream applications. In fact, an attempt at wide-range cell-free expression systems has been developed that combine cell lysates from 10 diverse bacterial species (53).

While the vast majority of the CFPS systems reported are based on cellular extracts, cell-free expression systems based on a mixture of tRNAs, ribosomes and pure, recombinantly expressed, histidine (His)-tagged components for transcription, translation and energy generation have been reported for *E. coli* (54) and *Thermus thermophilus* (55). The reconstituted *T. thermophilus* system is not commercially available, but the highly optimized PURE (Protein synthesis Using Recombinant Elements) system from *E. coli* has been commercialized as NEB PURExpress®. Reconstituted systems have the advantage of being protease- and nuclease-free compared to traditional S30 cell extracts where all soluble cytosolic components are present in the extract. Thus, linear nucleic acids, such as PCR products and mRNAs, and the synthesized proteins are more stable in PURE systems. Furthermore, an additional advantage of such systems is the absence of competing side reactions such as nonspecific phosphatases (54), which rapidly degrade the phosphorylated molecules used as energy sources. However, important cofactors or chaperones may be missing in these purified systems, and lead to inefficient folding of the target protein, which together with its high cost represent the major disadvantages of the PURE system. In contrast, crude cell extracts represent an inexpensive route to cell-free protein synthesis, reaching recombinant protein yields of up to 2.34 mg/ml (56,57), which makes extract-based CFPS a better candidate than the PURE system for scalability into high-volume fermentation conditions (31,56).

The majority of CFPS systems available are active at moderate temperatures (20–40 °C) but there are several reasons to explore the possibilities of protein synthesis at higher temperatures, such as enabling the functional expression of highly thermostable recombinant proteins that do not fold properly at room temperature and reducing mRNA secondary structures that otherwise might be inhibitory for translation (58). However, a disadvantage of CFPS operating at higher temperatures is the accelerated degradation of template nucleic acids. *In vitro* translation extracts from different thermophilic bacteria and archaea have been reported since 1993. Ruggero *et al*. reported an efficient *in vitro* translation (IVT) of archaeal natural mRNAs at high temperature (75 °C), with extracts of the extreme thermophile *Sulfolobus solfataricus* (59). Endoh and coworkers reported another system for cell-free protein system combining extracts from the hyperthermophilic archaeon *Thermococcus kodakarensis* KOD1 with a thermostable T7 RNA polymerase. Protein synthesis at high temperatures has also been demonstrated *in vitro* for thermophilic bacteria: Ohno-Iwashita *et al*., 1975 reported the synthesis of poly(Phe) with a cell-free extract of *T. thermophilus* at 65 °C (60) and more recently Zhou and coworkers described thermostable IVT of superfolder GFP (sGFP) using purified thermostable components (55).

The absence of a physical boundary during CFPS creates a significant challenge in the application of *in vitro* transcription and translation (IVTT) to analyze individual gene variants from metagenomic or protein variant libraries for enzyme discovery and engineering, respectively. The compartmentalization of the IVTT reaction is strictly needed to establish a linkage between the observed (and selected) phenotype and its encoding genotype. Furthermore, given the high cost of IVTT extracts and the throughput required to interrogate at least a fraction of the natural or artificial diversity contained in (meta)genomic libraries, even the use of the smallest microwell plates would compromise the economic viability of the application of IVTT to high-throughput screening. Nevertheless, compartmentalizing IVTT systems within monodisperse aqueous droplets with cell-like diameters (2-200 μm) and femto- to nanoliter volumes dispersed in an immiscible perfluorocarbon, may unlock the economic feasibility of *in vitro* screenings. In fact, large numbers of droplets (∼10^7^ -10^9^ in one experiment) can be produced at a more reduced cost per assay (∼10^6^ -fold) (61) than industrial robotic screening platforms. Amongst the successful experiments in directed evolution that involve *in vitro* expression in microdroplets are selections of enzymes such as DNA methyltransferases, phosphotriesterases, or glycosidases (62–64) in polydisperse droplets and more recently, proteases using monodisperse droplets created with microfluidics (65). Since the success of such directed evolution experiments in droplets is dependent on the efficiency of protein production in droplets by IVTT, *in vitro* expression can be boosted by generating multiple copies of the DNA template after emulsion PCR (66) or rolling circle amplification (RCA). To that end, two solutions have been proposed: either a two-step workflow with amplification of encapsulated single DNA molecules on beads (67) or a three-step workflow that makes use of picoinjection to sequentially add IVTT components and a fluorogenic substrate to microfluidic droplets containing the product of RCA (65).

Considering the relevance of thermostability as a property for industrial enzymes to enhance operational stability, this work aimed to develop thermostable cell-free protein synthesis of thermozymes that: i) enables a one-pot enzymatic assay; and ii) is compatible with droplet microfluidics. To the best of our knowledge, this is the first description of enzyme assays powered by IVTT at high temperature, compatible with microfluidic droplets, which will undoubtedly provide a cost-effective, simple and powerful tool for ultrahigh-throughput screening of libraries for enzyme discovery and evolution as well as open more opportunities in different applications from protein biochemistry to biomedical science in extremophile research.

## METHODS

### Strains and growth media

*E. coli* was grown at 37 °C in Luria-Bertani lysogeny broth (LB; 10 g/L Tryptone, 10 g/L NaCl and 5 g/L yeast extract) and *T. thermophilus* was grown at 65 °C in Thermus Broth (TB; 8 g/L Tryptone, 4 g/L NaCl and 3 g/L yeast extract in carbonate-rich mineral water). Media were solidified by the addition of 2% (w/v) of agar, if needed, and/or supplemented with a final concentration of 30 µg/ml kanamycin (Kan), 100 µg/ml ampicillin (Amp) or 20 µg/ml chloramphenicol (Cam) for selection.

*E. coli* DH5*α* [*sup*E44, Δ*lac*U169 (φ80 *lacZ*ΔM15), *hsd*R17, *recA*, *end*A1, *gyr*A96, *thi*^-1^ *relA1*] was used for construction of plasmids, whereas electrocompetent *E. coli* BL21(DE3) [*hsd*S, *gal* (λcIts857, *ind*1, *Sam7*, *min5*, *lacUV5*-T7 gene 1] and *E. coli* BL21 Rosetta [F^-^ *ompT hsdS_B_* (r ^-^m_B-_) *gal dcm* (DE3) pRARE (Cam^R^)] were used for overexpression and purification of recombinant proteins.

*T. thermophilus* strain HB27 was used to develop and optimize the IVTT protocol and the composition of CFPS reaction mixes. To reduce the background activity when coupling IVTT with enzymatic activity, S30 extracts were generated with *T. thermophilus* strain BL03, deficient in major hydrolytic activities including the gene encoding glycosidase *TTP0042*(68).

### Nucleic acid manipulation and transformation

Isolation of plasmid DNA was carried out with Genejet Plasmid Miniprep kit (Thermo Fisher Scientific, Pittsburgh, PA, USA), according to the manufactureŕs instructions. DNA was amplified by polymerase chain reaction (PCR), using GoTaq Flexi Polymerase (Promega, Madison, WI, USA), while for high fidelity amplification, PfuUltra II HS polymerase (Agilent Genomics, Santa Clara, CA, USA) was employed, following the manufacturer’s protocol. DNA fragments such as PCR products or DNA fragment as an agarose gel slices were purified using the Wizard® SV Gel and PCR Clean-Up System kit from Promega (Madison, WI, USA). DNA was digested with the appropriate restriction endonucleases (FastDigest, Thermo Fisher Scientific, Pittsburgh, PA, USA) following the manufacturer’s instructions. To avoid religation, vector DNA was dephosphorylated using FastAP Thermosensitive Alkaline Phosphatase (ThermoFisher Scientific, Pittsburgh, PA, USA). For routine plasmid construction, digested vectors and insert DNA fragments were ligated using T4 DNA Ligase (Promega, Madison, WI, USA) following manufacturer’s recommendations. DNA concentration was measured with Nanodrop™ One (Thermo Fisher Scientific, Pittsburgh, PA, USA). Sanger DNA sequencing was performed by Macrogen Inc. (Seoul, Republic of Korea) and DNA sequences were analyzed using SnapGene software (GSL Biotech, Boston, MA, USA).

The genes encoding pyruvate kinase (PK, gene name *TT_C1611,* UniProt ID: Q72H84), nucleoside diphosphate kinase (NDK, *TT_C1798,* Q72GQ0), adenylate kinase (ADK, *TT_C1307,* Q72125), lactate dehydrogenase (LDH, *TT_C0748,* P62055) and inorganic pyrophosphatase (IPP, *TT_C1600,* Q72H95) were amplified by PCR from *T. thermophilus* HB27 genomic DNA extracted using the DNeasy UltraClean MicrobialKit (QIAGEN). Amplified genes were digested and inserted into the pET28b(+) vector (Merck Millipore) (NDK, ADK, PK, IPP) or pET28b(+) vector (Merck Millipore) (LDH). LDH, PK, NDK and IPP were cloned with NdeI and HindIII restriction enzymes (ThermoFisher Scientific, Pittsburgh, PA, USA), whereas ADK was cloned with NdeI and EcoRI (ThermoFisher Scientific, Pittsburgh, PA, USA). Chemically competent *E. coli* DH5α cells were transformed with the constructs. Transformation was carried out by heat shock following the method described by Hanahan (69). Five microliters of ligation product or 100-200 ng of plasmid was added to a 50 µl of competent cells, which were further incubated on ice for 30 min. Then, the tubes were heated at 42 °C for 90 s. After cooling down on ice for 5 min, 350 µl of SOC medium was added and incubated in a shaker at 37 °C for 1 h for the expression of ampicillin or 3 h for the expression of kanamycin resistance.

Clones were checked by restriction digestion with the corresponding enzymes, followed by sequencing. To ensure correct in-frame expression of the His_6_-tag-encoding sequence, ADK required additional site-directed mutagenesis, performed using QuikChange II XL Site-Directed Mutagenesis Kit (Agilent Technologies, Santa Clara, CA, USA) followed by transformation of *E. coli* DH5α cells and confirmative sequencing.

The gene encoding superfolder GFP (sGFP) was amplified by PCR using plasmids pET28b (+)_sGFP as template (Supplementary Table 1) and the T7 primers indicated in Supplementary Table 2.

### Solutions and chemicals

All IVTT buffers and solutions were prepared with diethylpyrocarbonate (DEPC)-treated water (0,05%) or nuclease-free water (Invitrogen, CA, USA). The S30A and S30B buffers along with the HEPES, nucleotide mix, potassium glutamate, magnesium glutamate, folinic acid and spermine solutions were prepared following the protocol described in Sun et al., 2013 (70). The amino acid solution was prepared according to Cashera & Noireaux, 2015 (71).

Chemicals were purchased in analytical grade from Merck KGaA (Darmstadt, Germany), Bio-Rad Laboratories (Hercules, CA, USA), or Sigma-Aldrich (St. Louis, MO, USA). Sigma-Aldrich synthesized the oligonucleotides used.

### Protein expression and purification

Electrocompetent *E. coli* BL21 (for ADK, PK, NDK and IPP) and *E. coli* BL21 Rosetta (for LDH) cells were transformed with the previously described constructs. Electroporation was carried out by mixing 45 µl of competent cells with 100-200 ng of plasmid and subjecting the cells to a short 5 ms electric pulse under a 12500 V/cm electric field in a Gene Pulser II® (Bio-Rad, Hercules, CA, USA) (2500 v, 201 Ω and 25 µF) using 0.2 cm gap cuvettes (Bio-Rad Gene Pulser). Immediately after the pulse, 500 µl of SOC medium as added and incubated at 37 °C during the appropriate time for each antibiotic resistance before plating on selective medium.

Transformant colonies were grown overnight in 20 mL LB medium supplemented with ampicillin (pET22b constructs) or kanamycin (pET28b constructs). Chloramphenicol was supplemented when using *E. coli* BL21 Rosetta as a host. Cultures were inoculated at a 1/100 dilution of those preinocula and induced at OD_600_ 0.5 with 0.5 mM IPTG for PK, IPP, NDK and ADK. LDH-transformants were induced at OD_600_ 0.7 and 1 mM of IPTG. Cells were grown at 37 °C for 5 h after induction (PK, PPI, NDK) or 20 h at 22 °C (ADK). *E. coli* BL21 Rosetta transformed with LDH was grown for 36 h at 17 °C. Cells were harvested and pelleted after induction and growth. Pellets were frozen and stored at -20 °C until further use. Negative controls were carried out with transformants harboring the corresponding empty vectors.

To verify the expression of proteins, pellets from different times after induction were resuspended in 50 mM phosphate buffer, sonicated (0.6 Amplitude and 50% pulse) and centrifuged at 14100 ×g for 10 minutes. The supernatant was separated and the pellet was washed with phosphate buffer 50 mM pH 7.5 and 0.1% w/v Triton X-100, analyzed by SDS-PAGE in a 12% acrylamide gel according to the method described by Laemmli (72). The gels were stained with Coomassie Brilliant Blue G-250 from Bio-Rad Laboratories (Hercules, USA).

To purify the protein pellets from ADK, NDK, PK and IPP (large scale production) were resuspended in 50 mM phosphate buffer, sonicated (0.6 Amplitude and 50% frequency) and centrifuged at 15000 ×g for 30 min. Proteins found in the soluble fraction were later purified by immobilized metal ion affinity chromatography (IMAC), using Talon resin (BD ClonTech). Purified proteins were concentrated using 10 kDa cutoff Amicon Ultra Centrifugal Filters (Merck Millipore) and stored in 50% v/v glycerol at -20 °C until further use.

After protein concentration, purity was checked by sodium dodecyl sulfate polyacrylamide gel electrophoresis (SDS-PAGE) in a 12% polyacrylamide gel using the Bio-Rad Protein Assay (Bio-Rad, Hercules, CA, USA), according to the manufacturer’s protocol, using bovine serum albumin (BSA) as standard.

### Western blot

Western blot analysis was carried out by semidry electroblotting on immobilon-P transfer membrane from Merck KGaA (Darmstadt, Germany). Anti-GFP polyclonal antibody and secondary goat-anti-rabbit HRP conjugate antibody were supplied by Thermo Scientific (Ulm, Germany) and used at the dilutions recommended by the manufacturer. The antibodies were diluted according to manufactureŕs recommendations. Detection of proteins was achieved by chemiluminescent reaction and capture on a film, using the charge-coupled device “X-OMAT 2000 Processor” from Kodak (Rochester, USA). The reaction was enabled by addition of a solution containing 100 mM Tris-HCl pH = 7.8, luciferin and luminol and a second solution containing 100 mM Tris-HCl pH 7.8 and H_2_O_2_ (enhanced chemiluminescent reaction). Quantification of sGFP was carried out by image analysis using Fiji image processing software (73) by comparison of lanes containing 8 μl of completed IVTT reaction with lanes containing 300 or 400 ng of pure sGFP.

### Activity assays of the energy regeneration enzymes

Enzymatic assays were performed at 60 °C. Four replicates of each sample in the endpoint assays and three replicates in the real-time assays were measured to assure statistical significance. Negative controls were performed by omitting the substrate. Spontaneous conversion without the enzyme was also performed as a negative control.

LDH was assayed at 60 °C in 50 mM Tris-HCl buffer, 0.2 mM NADH and 0.3 mM pyruvate. The assay was performed at pH values ranging from 5.5 to 8 and at different ionic strength values ranging from 4 to 40 mM (74). One unit of LDH activity was defined as the amount of enzyme able to convert 1 micromole of pyruvate per minute.

PK was assayed with 5 mM phosphoenolpyruvate (PEP), 5 mM ADP, 20 mM KCl and 5.4 mM MgSO_4_ in 100 mM Tris buffer pH 7.5. In this case, PK was assayed using a coupled assay with LDH and recording NADH absorbance at 340 nm with 5 mM ADP, 40 mM KCl, 40 mM MgSO_4_, 6 mM PEP, 0.4 mM NADH and 6.3 mU of LDH in 50 mM Tris buffer pH 7 (75). One unit of PK activity was defined as the amount of enzyme able to convert 1 micromole of PEP per minute. NDK was assayed with 40 mM KCl, 40 mM MgSO4, 6.3 mU LDH, 7.06 mU PK, 1 mM GDP, 0.2 mM NADH, 1.1 mM PEP and 2.2 mM ATP in 50 mM Tris-HCl buffer pH 7. One unit of DNK activity was defined as the amount of enzyme able to convert 1 micromole of GDP per minute.

ADK was assayed with 2.5 mM ADP, 100 mM KCl and 2 mM MgCl_2_ in 50 mM Tris-HCl buffer pH 7.5. Different dilutions of the enzyme preparation were assayed for 1 hour and analyzed with a luciferin-luciferase coupled reaction (76) using the CLSII-Bioluminiscent ATP Assay Kit (Roche).

IPP was assayed with 5 mM pyrophosphate (PPi), 120 mM NaCl, 5 mM KCl and 5 mM MgCl_2_ in 20 mM Tris-HCl buffer pH 7.5. For the end-point assay different dilutions of the enzyme were assayed and stopped at different times (77). The released phosphate was analyzed with the malachite-green phosphomolybdate assay by adding 200 μl of the malachite green reagent (5.72% w/v ammonium molybdate in HCl 6M, 2.32% w/v PVA, 0.0812% w/v malachite green; 1:1:2:2) and measured at 640 nm (78).

### In vitro transcription and translation extracts

Preparation of S30 extracts was performed by a modification of the method published by Sun *et al.* (70) under RNase-free conditions. Briefly, a 30 ml preculture of *T. thermophilus* HB27 was grown overnight aerobically under rotational shaking (150 rpm) at 65 °C in liquid TB (79). The preculture was used to inoculate 1.2 L culture with TB medium which was cultured under aerobic conditions at 65 °C until A600 reached 1.0–1.2 (corresponding to the mid-log growth phase). Immediately after growth the culture was cooled down to 4 °C. Cells were harvested by centrifugation at 5000 ×g for 15 min at 4 °C and washed with 500 mL of DEPC-treated water (0.05%). The cell pellet was resuspended in 250 mL of buffer S30A (14 mM Mg-glutamate, 60 mM K-glutamate, 50 mM Tris, pH 7.7, 2 mM DTT) and centrifuged once more at 5000 ×g for 15 min. Fifty milliliters of S30A buffer was used to wash the cells one last time. The pellet was weighed before being stored at –80 °C. The cell pellet was transferred into a prechilled mortar and disrupted by grinding with 1.5 g of precooled alumina (Sigma‒Aldrich, St. Louis, MO, USA) per gram of wet cell for 15 min over ice. The cell slurry was resuspended in buffer S30A (0.5 mL for gram of wet cell) and transferred to a centrifuge tube. The extract was centrifuged at 30000 ×g for 30 min at 4 °C to remove alumina and cell debris. The resulting supernatant was incubated at 37 °C for 80 min with shaking at 200 rpm and then centrifuged at 12000 ×g for 10 min at 4 °C. The S30 extract was dialyzed against S30B buffer (14 mM Mg-glutamate, 60 mM K-glutamate, ∼5 mM Tris, pH 8.2, 1 mM DTT) in 3k MWCO dialysis cassettes (Thermo Fisher Scientific, Pittsburgh, PA, USA) for 3 h at 4 °C. The extract concentration was determined by Bradford (Bio-Rad, Hercules, CA, USA). Aliquots of the S30 cell extract were then frozen in dry ice and stored at – 80 °C.

### In vitro transcription and translation in bulk

The CFPS reaction was performed in a final volume of 25 µl using plasmid DNA as a template, unless stated otherwise. The reaction mixture contained the S30 extracts and the IVTT buffer mix with the ingredients shown in Table 1 and a thermostable T7 RNA polymerase (Toyobo). RNase inhibitor was purchased from New England Biolabs (Ipswich, MA). ATP, GTP, UTP and CTP were purchased from Promega (Madison, WI). Total tRNAs from *E. coli* were purchased from Sigma-Aldrich (St. Louis, MO).

**Table 1.**
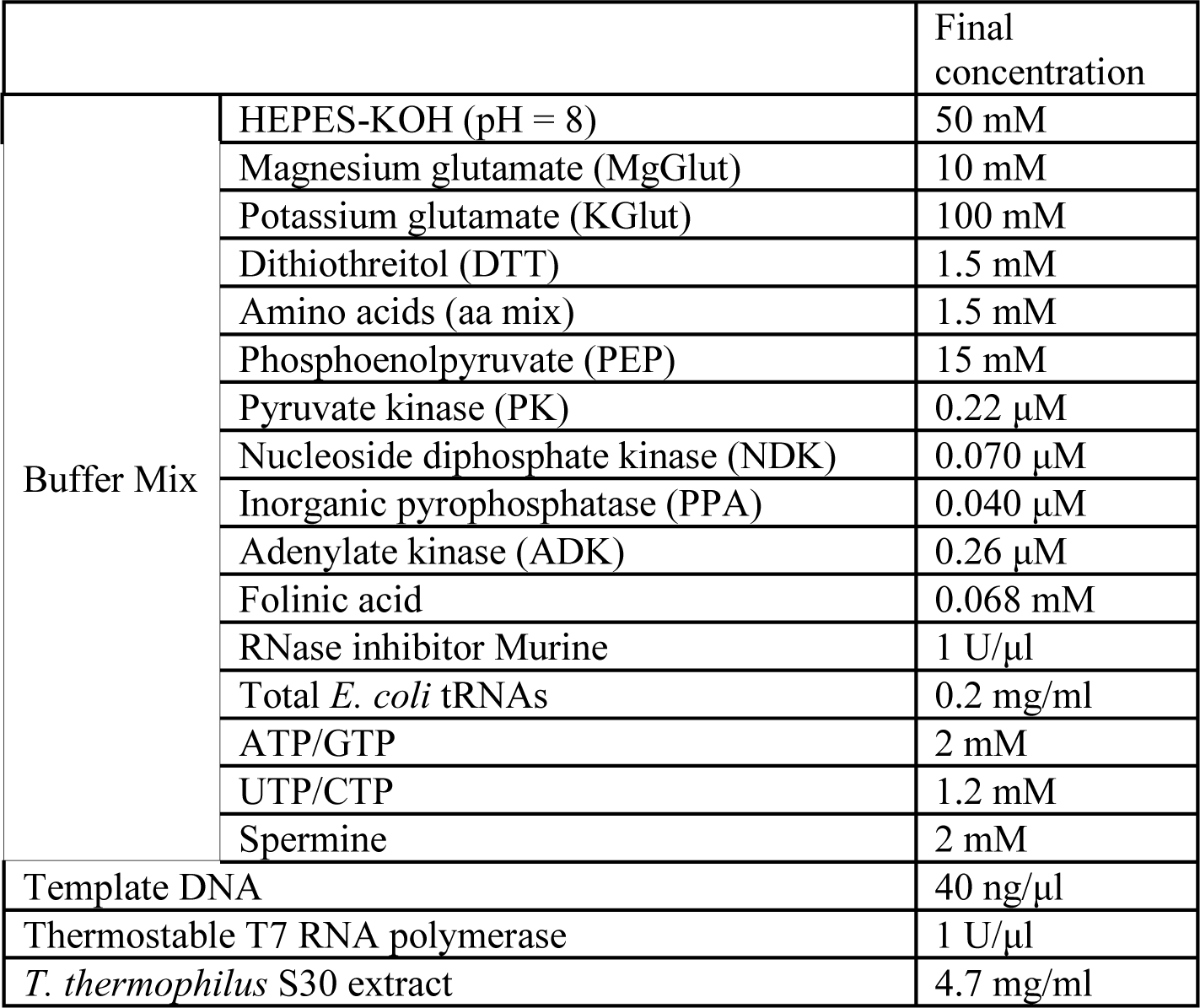
Composition of the reaction mix for thermostable IVTT.

When the IVTT was coupled to an enzymatic activity assay, a fluorogenic substrate was added to the medium at a final concentration of 5 µM from a 10 mM stock prepared by dissolving substrates in DMSO. The substrates used were fluorescein di-β-D-glucopyranoside (FDGlu), fluorescein di-β-D-cellobioside (FDC) purchased from Thermo Fisher Scientific and AAT Bioquest, respectively and fluorescein di-beta-galactoside and esterase substrates fluorescein dicaproate (FD6) and fluorescein dilaurate (FDL) purchased from abcr GmbH and Merck, respectively. The samples were transferred to a real-time PCR cycler (Rotorgene 6000, Corbett Research) and incubated at the indicated temperatures for the indicated time. Fluorescence was measured in the high-resolution melt (HRM) channel (excitation 460±20 nm, emission 510±5 nm) and the gain was adjusted manually according to the fluorescence reading during the first ten cycles of the reaction.

### In vitro transcription and translation at different temperatures

Cell-free transcription and translation reactions were carried out using PURExpress® *In Vitro* Protein Synthesis Kit (NEB) following the manufactureŕs protocol, adjusted to 10-µl reaction volume and using 5 ng of pET28_*sGFP* as a template. Fluorescence was measured using Rotorgene 6000 (Corbett Research) in the high-resolution melt (HRM) channel (excitation 460±20 nm, emission 510±5 nm) and the gain was adjusted manually according to the fluorescence reading during the first ten cycles of the reaction.

### Rolling Circle Amplification (RCA) coupled with IVTT

Plasmid DNA was amplified using REPLI-g® Midi Kit (QIAGEN), according to the manufactureŕs instructions. One nanogram of template plasmid was used in 25 µl final volume and the reactions were incubated isothermally at 30 °C for 3 h, followed by inactivation at 65 °C for 3 min. Then, components for IVTT were added to the reaction, either *T. thermophilus* extracts or PURExpress® *In Vitro* Protein Synthesis Kit (New England Biolabs, Ipswich, MA, USA) and the reactions were further incubated at either 50 °C or 37 °C for another 4 h in the case of *T. thermophilus* extracts or 2 h for the *E. coli* PURE system.

### In vitro transcription and translation in droplets

Designs of flow focusing chips used for droplet generation were obtained from DropBase (https://openwetware.org/wiki/DropBase:Devices). Device master molds were microfabricated by Tekniker (Eibar, Spain) on silicon wafers by soft photolithography with a 30 µm height. A mixture of poly(dimethylsiloxane) (PDMS, Sylgard 184 Dow Corning (Midland, USA) and cross linker (ratio 10:1 w/w) was poured over the master mold, then degassed and then cured overnight at 65 °C. The cured device was cut and peeled from the master, and holes for tubing were cut with a 1-mm biopsy punch (Kai Medical, Solingen, Germany). After treatment with oxygen plasma for 15 seconds (Diener Femto, 30 W, 40 kHz), the device was sealed against a glass slide. The channels were washed with a 1% (v/v) solution of trichloro(1H,1H,2H,2H-perfluorooctyl)silane in HFE 7500 and baked for at least 3 h at 65 °C.

The flow was driven with Nemesys Base 120 module and Nemesys S pumps (Cetoni GmbH (Korbußen, Germany) syringe infusion pumps using 1 mL and 100 μL Gastight syringes, (Hamilton, Reno, USA) connected to fine-bore polyethylene tubing with 1.09 mm outer diameter and 0.5 mm inner diameter (Smiths Medical). We tested several combinations of surfactants Pico-Surf (Sphere Fluidics) or RAN 008 Fluorosurfactant (RAN Biotechnologies) with fluorocarbon oils HFE7500, FC70 or FC40 (Merck Millipore) as the continuous phase. The microfluidic equipment was integrated by an inverted microscope (Leica DMi8) connected to a high-speed camera (Fastcam Mini UX 50, Photron) for real-time visualization of experiments. Emulsions were routinely photographed in Fast-Read 102 slides with counting chambers (Biosigma s.r.l., Italy) using an Olympus BX50 microscope equipped with a Pike F-032B camera (Allied Vision Technologies) and a 25x objective. Dimensions were determined from at least n=40 droplets using Fiji image processing software(73).

## RESULTS

### Preparation of IVTT extracts from Thermus thermophilus HB27

The preparation method of cell lysates for cell-free protein synthesis is a crucial step for their functionality, largely affecting the yield of synthesized protein (35,80,81). The so-called S30 extracts for CFPS are composed of a soluble cell fraction that contains all the cytosolic enzymes required for transcription and translation. For the preparation of S30 extracts from *T. thermophilus*, we modified an existing protocol for *E. coli* (70) under RNase-free conditions. The resulting protocol involved rapid growth and harvesting of *T. thermophilus* cells at exponential phase when intracellular translation is at its peak, washing the cells, mechanical lysis cell lysis by manual grinding using a prechilled mortar and a pestle in the presence of pre-cooled alumina instead of a bead beater and glass beads and activation of the extract through a run-off reaction, a process believed to degrade endogenous mRNA transcripts and genomic DNA that can produce unwanted side products and reduce cell-free translation efficiency. Furthermore, we added a dialysis step to remove inhibitory small molecules (less than 10 kDa) that affect the final protein yield.

Then, supplements were considered in order to improve protein folding, stability and yield. First, dithiothreitol (DTT) was added to the cell-free system to increase protein expression and to preserve the reducing character of the cytoplasmic environment (82). DTT has little effect on the folding of cytoplasmic proteins and a significantly greater effect on the folding of proteins that require the formation of disulfide bonds for activity (83). Additionally, spermine was added, instead of spermidine, since it has been shown that peptide synthesis occurs most rapidly when spermine is added to the cell-free system medium reaction (84). In addition, tRNAs from *E. coli* (Sigma) were added to the mix. Finally, to achieve transcription at higher temperatures, a thermostable T7 RNA polymerase (tT7 RNApol), with an optimum reaction temperature of approx. 50 °C, was incorporated into the mix.

Finally, the supply of energy (ATP and GTP) has been reported to be a limiting factor in cell-free protein synthesis (85,86). The simplest method is to add high energy compounds to fuel IVTT, such as phosphoenolpyruvate (PEP)(87), 3-phosphoglycerate (3PGA) (88) or creatine phosphate (CP) (34,86,89) and convert them into ATP and/or GTP *via* an enzymatic cascade. To achieve the energy regeneration at high temperature and improve the productivity of our system, we cloned, expressed, purified and characterized the thermostable enzymes from *T. thermophilus* involved in two energy-regeneration systems. The multienzymatic ATP-recycling cascade is composed of pyruvate kinase (PK), adenylate kinase (ADK) and inorganic pyrophosphatase (IPP). The GTP-recycling cascade is composed of pyruvate kinase (PK) and nucleoside diphosphate kinase (NDK). These systems use phosphoenolpyruvate as phosphate donor and provide constant ATP for the aminoacylation of tRNAs and GTP to support translation initiation and elongation. The individual activity and stability of each added enzyme were determined and are shown in Supplementary Table 3.

### Coupled transcription and translation at high temperature

We generated S30 extracts from *T. thermophilus* cells, depleted endogenous DNA and RNA and supplemented them with free amino acids and tRNAs as described above. Protein expression was initiated by the addition of a suitable template (pET28sGFP). We chose the superfolder GFP (sGFP) as the target protein since it can be expressed and it is able to properly fold and fluoresce when expressed in *T. thermophilus* at 70 °C (90). Moreover, sGFP synthesis can be easily detected and measured in real time by its fluorescent emission.

To determine the optimal type of DNA template to use in our cell-free expression system, different types of DNA template were added to the CFPS extracts and incubated at 50 °C for 120 min and sGFP expression levels were compared (Figure 1A). We tested the 957 bp amplicon of the sGFP gene (PCR sGFP, linear DNA), a pET28b(+) plasmid expressing sGFP (pET28b_sGFP, circular DNA) under the control of the strong φ10 promoter for the T7 RNA polymerase (T7 promoter), and an empty pET28b(+) vector as negative control of the reaction. Upon mixture and incubation of all components in a real-time thermocycler, a rapid accumulation of protein product was observed after 20 min and reached saturation at 40 min when circular DNA was used as template (Figure 1A). However, we observed an approximately 3-fold lower amount of synthesized protein when linear DNA was used as a template. We did not detect any fluorescence when the empty vector was used as a template, as expected. From this result, we can conclude that extracts are functional and that circular DNA templates are preferred, likely due to the degradation of linear DNA by exonucleases naturally present in the extracts.

**Figure 1.**
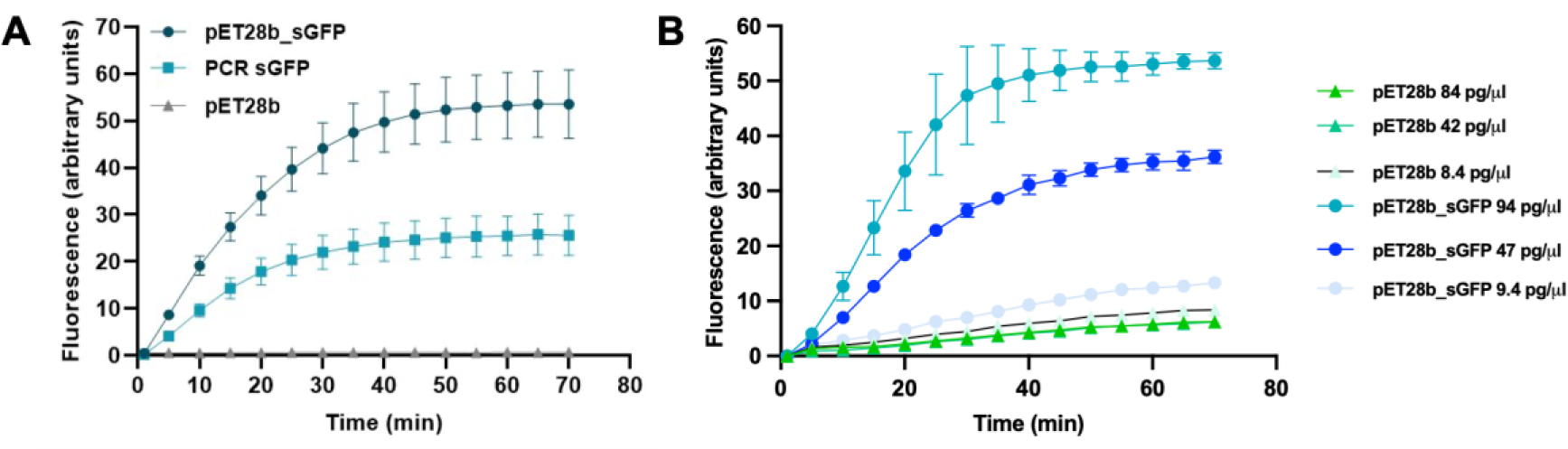
Synthesis of sGFP using *T. themophilus* S30 extracts and different types of DNA template. **(A)** Reaction mixtures containing 40 ng/µl of pET28b_*sGFP* (circles) or the same molar concentration of sGFP PCR amplicon (squares) were incubated at 50 °C for 70 min. As a negative control of the reaction, an empty pET28b vector was used (triangles). Composition of the reaction mixtures are indicated in Table 1. sGFP synthesized was monitored in real time as fluorescence emission. (**B**) Different amounts of template DNA (in ng/ml) were tested as in (A), with the corresponding amounts of empty vector as negative controls. Results are the average of n=3 reactions and error bars represent standard deviations.

Then, we sought to determine the sensitivity of our system by using a decreasing number of DNA template molecules of in the reaction. We tested several amounts of pET28b_sGFP DNA (84, 42 and 8.4 pg/μl) and the same number of molecules of empty pET28b(+) (amounting to 94, 47 and 9.4 pg/μl), as negative control. We were able to detect fluorescence in the presence of DNA amounts as low as 9.4 pg/μl using a real-time thermocycler as the detection platform. A minimal fluorescence signal was detected when an empty plasmid was used in the reaction (Figure 1B), likely due to the experiment being performed at the maximum instrument gain.

We also determined the temperature limit of our *in vitro* transcription and translation-coupled protein synthesis system. To that end, the reaction mixture using pET28b_sGFP as template was incubated at different temperatures ranging from 37 °C to 60 °C, above and below the optimum for the tT7RNApol, for 3 h. Maximum levels of protein synthesis were observed at 50 °C and some synthesis was even detected at 37 °C (Figure 2). However, no significant synthesis of sGFP was observed when the reactions were carried out at or above 55 °C. From this result, it appears that the optimum temperature for our reaction is 50 °C, likely due to the limited stability of the tT7 RNA polymerase compared to the rest of the components.

**Figure 2.**
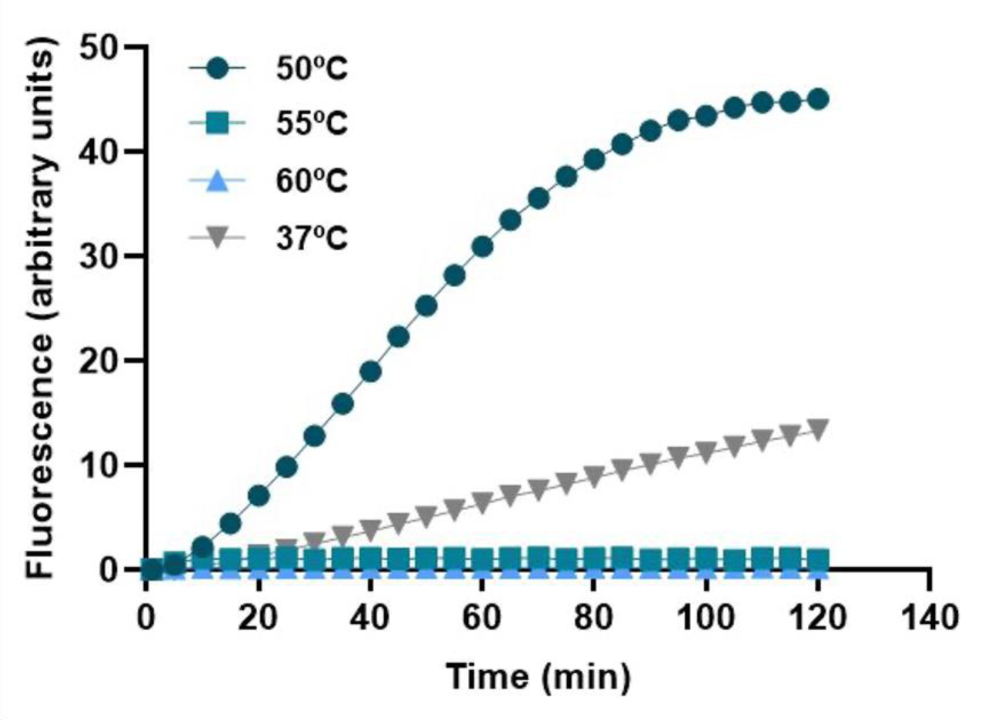
Temperature limit of sGFP synthesis with *T. thermophilus* S30 extracts. Reaction mixtures containing 40 ng/µl of pET28b_*sGFP* were incubated at 37 °C (inverted triangles), 50 °C (circles), 55 °C (squares) and 60 °C (triangles) for 120 min. Composition of the reaction mixtures are indicated in Table 1. sGFP synthesized was monitored in real time as fluorescence emission.

Given the fact that increasing the temperature above 50 °C was not conducive to increasing the yield, we tried to increase the amount of tT7 RNA polymerase instead. When sGFP synthesis was carried out in the presence of different amounts of tT7 RNA polymerase (between 0.5 and 2.5U/µl) and incubated at 50 °C for 2 h, we did not observe a linear increase in protein yield suggesting that tT7RNApol may not be the rate-limiting factor in the CFPS reaction mix (Supplementary Figure 1).

The amount of synthesized sGFP after completion of a CFPS reaction at the optimum temperature of 50 °C was quantitated both by image analysis of a Western blot comparing with known amounts of sGFP (Figure 3A) and by interpolating the fluorescent signal registered in a real-time thermocycler with a sGFP calibration curve acquired with the same parameters (Figures 3B-D). The former method yielded 67.7 ng/µl sGFP whereas the latter yielded 56.4 ng/µl from the same CFPS reactions. Other CFPS reactions carried out and acquired under the same conditions yielded 75.7 ng/µl (Figure 1) and 64.7 ng/µl (Figure 2), which are in a similar range.

**Figure 3.**
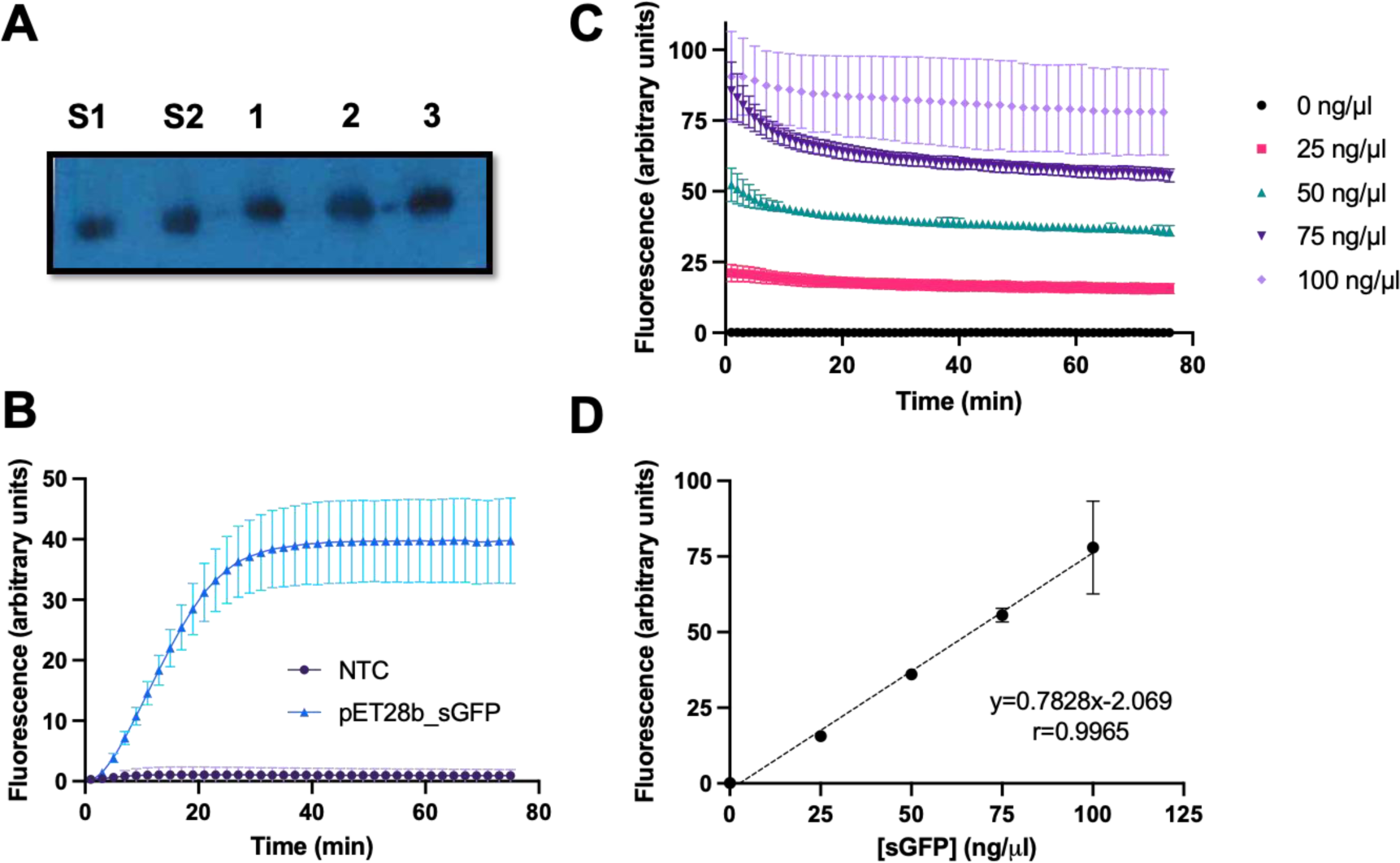
Quantification of the yield of CFPS reactions. Reaction mixtures containing 40 ng/µl of pET28b_*sGFP* as template were incubated at 50 °C for 70 min in a real-time thermocycler and the yield of sGFP was quantitated independently using both Western Blot and a calibration curve of sGFP. A: Western blot analysis of 8 μl aliquots of three independent reactions using an anti-GFP antibody and the ECL developing reaction, S1: 300 ng purified sGFP as standard, S2: 400 ng purified sGFP, 1-3: three replicate CFPS reactions. B: Average progress of the 3 independent CFPS reactions analyzed by Western Blot in panel A. C: Fluorescence of sGFP standards determined in triplicate after incubation for 70 min at 50 °C in a real-time thermocycler; D: calibration curve of sGFP standards after 70 min incubation at 50 °C. Error bars represent standard deviation.

### Coupling cell-free protein synthesis with DNA amplification

To further increase sensitivity, we sought to couple DNA amplification with IVTT. PCR-generated amplicons are often used as templates for CFPS but as shown above, they are not the optimal template for our system (Figure 1A). Since plasmids are the preferred template, we chose isothermal random and multiple-primed rolling circle amplification (RCA) using Φ29 DNA polymerase (91) to perform the amplification of template DNA. In our case, both operations must necessarily be performed sequentially due to the differences in the stability of the enzymes and components responsible for DNA amplification and IVTT.

To examine whether RCA products can serve as template for the *T. thermophilus*-based CFPS system, we selected the REPLI-g® Midi Kit (QIAGEN) for DNA amplification and *T. thermophilus* extracts described in this work or commercially available *E. coli* -based reconstituted protein synthesis system (PURExpress®, New England Biolabs) for the IVTT step using one ng of pET22b_sGFP as input DNA. To reduce the cross-inhibition between RCA and IVTT and in accordance to similar assays in the literature (65), we tested different RCA:IVTT volumetric ratios (1:1, 1:2 and 1:5). To determine the contribution of the input plasmid (1 ng) to the total protein synthesis yield, a control reaction was carried out without the addition of the RCA components.

When reactions were performed using the *E. coli* reconstituted system, significant levels of protein synthesis were detected after 30 min incubation, in particular when a 1:5 RCA:IVTT ratio was used. Under this condition, the reaction saturates very rapidly at 30 min incubation (Supp Fig 2A). However, no improvement in yield was obtained when IVTT was performed with *T. thermophilus* extracts at 50 °C (Supp Fig 2B). These results indicate that RCA and IVTT reactions cannot be coupled using *T. thermophilus*-based CFPS, even if the two reactions are carried out sequentially.

### Coupling cell-free protein synthesis with enzyme activity assays

Although we have shown that *T. thermophilus*-based CFPS allows the synthesis of proteins at high temperature (Figures 1 and 2), library screening for enzyme discovery or engineering requires carrying out an enzyme assay either simultaneously or after protein synthesis, preferably in a one-pot setup. Consequently, either fluorescein di-β-D-glucopyranoside (FDGlu) or fluorescein di-D-β-cellobioside (FDC) was added to the IVTT mix and the reaction was started by addition of the pET22b vector harboring a gene coding for a promiscuous glycosidase (*TTP0042*) from *T. thermophilus* HB27 (92) as template. Since the optimum temperature for *TTP0042* activity is higher than 50 °C (92), which is the limit for thermostable IVTT (Figure 2) reactions were first incubated at 50 °C for 2 h to allow protein synthesis and then, the temperature was increased to 70 °C for 17 h. As shown in Figure 4, we detected a prominent β–glucosidase activity in the presence of its specific substrate FDGlu and a low-level cellobiose hydrolase activity when the reaction was performed using FDC. We observed background activity when the reactions were performed using the empty plasmid as a template, likely due to autohydrolysis or the presence of other glycosidases in the S30 extracts. In fact, fluorescein dihexanoate, showed a very fast autohydrolysis, preventing activity determination of carboxyl esterases that are known to cleave this substrate (Supplementary Figure 3). Nevertheless, these results demonstrate that, given an adequate substrate, enzyme activity assays can be coupled to thermostable CFPS under the tested conditions.

**Figure 4.**
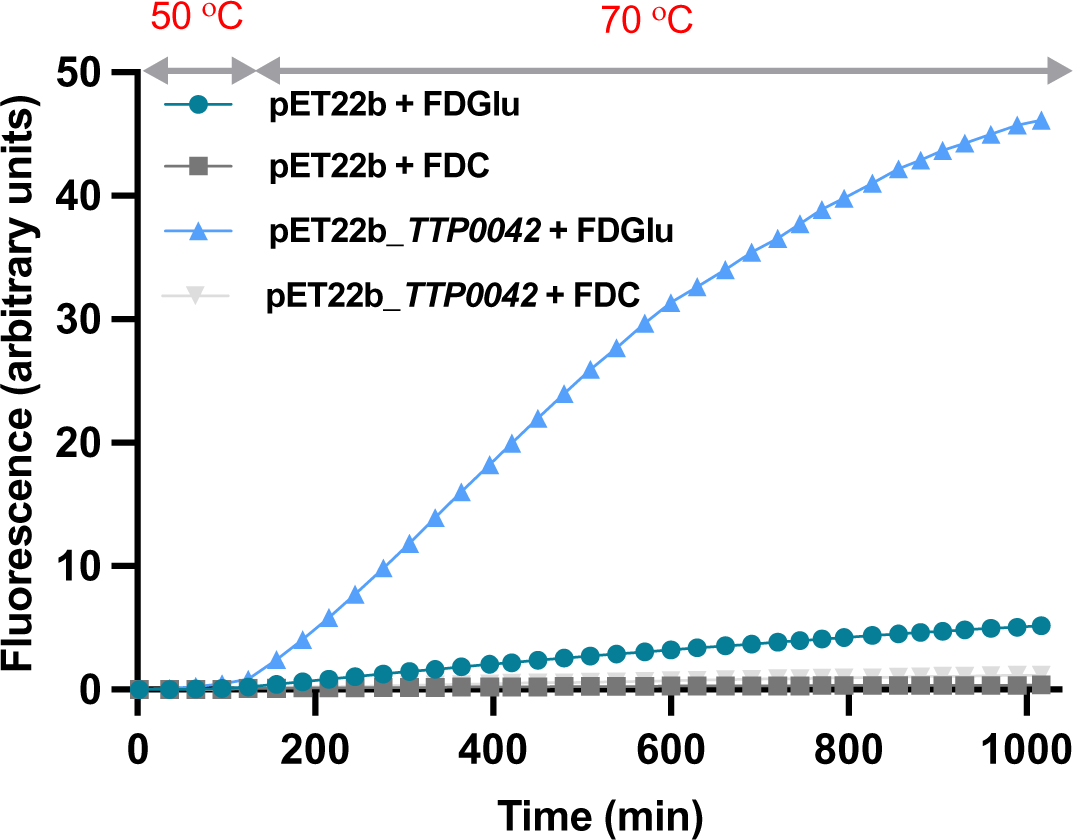
Coupling cell-free protein synthesis with an enzymatic activity. Reaction mixtures containing 40 ng/µl of pET22b_*TTP0042* and 5 µM of either FDGlu (triangles) or FDC (inverted triangles) were first incubated at 50 °C for 2 h and then, temperature was increased to 70 °C for 17 h. An empty pET22b plasmid was incubated in the presence of FDGlu (circles) or FDC (squares) as negative controls of the reactions. Composition of the reaction mixtures are indicated in Table 3. Results are the average of n=3 reactions and error bars represent standard deviations.

### Encapsulation in water-in-oil droplets

Cell-free extracts could be successfully encapsulated in water-in-oil droplets using PDMS flow focusing chips with standard designs. Several combinations of oils and surfactants were tested for compatibility with the viscosity, salt content and functionality of the IVTT extracts. The use of 1-1.5% RAN 008 fluorosurfactant in HFE 7500 provided the best droplet stability both during bpth flow focusing and incubation. As shown in Figure 5, droplets were monodisperse prior to and after incubation and fully compatible with a 24 h incubation at 50 °C, successfully achieving thermostable *in vitro* synthesis of sGFP in droplets for the first time, to the best of our knowledge.

**Figure 5.**
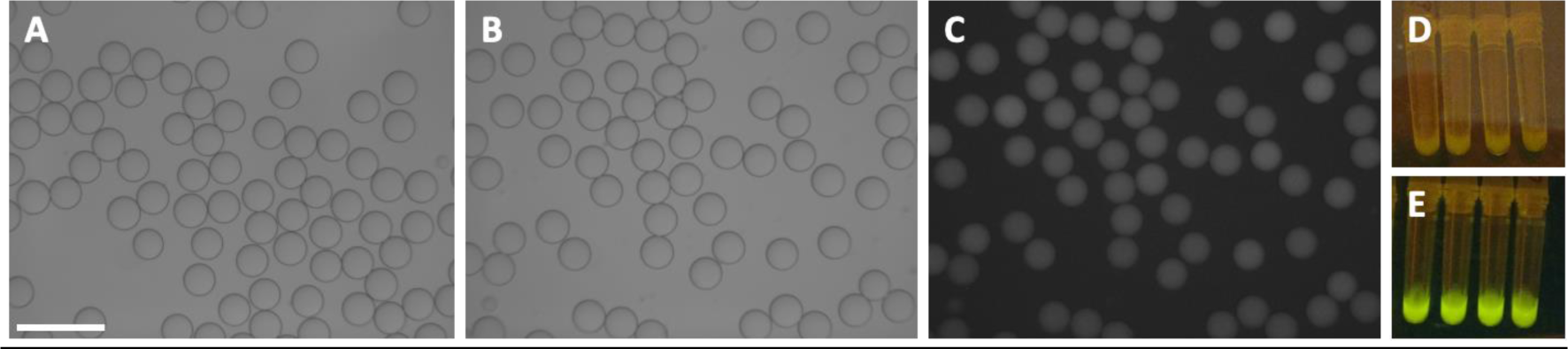
Cell-free protein synthesis in water-in-oil microfluidic droplets. Reaction mixtures containing 30 ng/µl of pET28b or pET28b_sGFP were encapsulated at 50 μl·h^-1^ using 30 μm flow focusing chips with 500 μl·h^-1^ 1% fluorosurfactant in HFE7500 as the continuous phase (A). The droplets, averaging 34 μm in diameter, were incubated for 3h at 50 °C in a real-time thermocycler and imaged in an Olympus epifluorescence microscope using brightfield illumination (B) or a FITC filter set (C). Cell-free protein synthesis with the pET28b template rendered emulsions (D) that were clearly distinguishable from emulsions with pET28b_sGFP (E) under blue light. Scale bar: 100 μm.

## DISCUSSION

In this study we describe a cell-free system for protein synthesis at high temperatures for the first time coupled to enzymatic assays and functional in microfluidic droplets. The use of cell extracts from thermophilic organisms provides several advantages, compared to mesophilic organisms due to the increase in temperature: the protein synthesis rate and substrate solubility are increased, folding of proteins from thermophiles is facilitated and the formation of mRNA secondary structures and the chances of microbial contamination are reduced (93).

Despite the fact that a reconstituted *Thermus thermophilus* translation system has been described and proven to be efficient at high temperatures (Zhou et al 2012), a major advantage of our S30 extract-based system is its simplicity when compared to the reconstituted system, which involves the overproduction and purification of 33 recombinant proteins, ribosomes and total tRNAs. Another reconstituted system from *E. coli* is commercially available (PURExpess®) providing a rapid method for *in vitro* protein expression (Shimizu et al 2001). The limitation of the latter is that it is not fully functional at any temperature above 37 °C (Supp Fig 4). The amount of synthesized protein using PURExpress® decreased by approximately 50% when *in vitro* transcription and translation were performed at 40 °C, compared to the amount obtained at 37 °C. No protein synthesis was detected at 45 °C or 50 °C in the Rotorgene 6000 (Supp Fig 4). Thus, our system offers a simple and rapid method able to give synthesized protein yields of approx. 70 ng/µl at elevated temperatures, providing a platform for thermostable protein expression and screening.

A general advantage of *in vitro* protein expression compared with host-based expression is the use of a PCR fragment as a DNA template in the reaction, as long as it contains a promoter and a ribosomal binding site (RBS) together with the translation initiation and termination signals. However, in this study, we observed a higher efficiency when using plasmid DNA as a template, compared to a PCR product (Figure 1). This is in agreement with the reported degradation of linear templates for cell-free expression by exonucleases naturally present in cellular extracts, primarily exonuclease V, encoded in the *recBCD* operon (94–96) and endonuclease I, encoded by *endA* (97). Using a linear DNA template would have advantages because it can be easily amplified by PCR, avoiding the time-consuming steps of cloning, plasmid amplification and extraction using bacterial cultures. Therefore, several alternatives can be implemented to increase the stability of linear templates and thus, increase the efficiency of cell-free protein synthesis, e.g.: obtaining IVTT extracts from *E. coli recBCD* knockout strains (98,99) or its equivalent addAB knockout mutants in *T. thermophilus* (100); from strains in which the genes coding for endonucleases V and I have been silenced before harvesting the cells (101); by adding additional exonuclease inhibitors to the reaction mix (88,102); depleting exonuclease V in the crude extracts (103); and protecting the ends of the linear dsDNA template by chemical modifications of its ends (104,105).

Nevertheless, using plasmid DNA as template and under optimal conditions, thermostable proteins such as sGFP or TTP0042 from *T. thermophilus* could be synthesized *in vitro* at 50 °C, reaching yields of 65 ng/ µl of sGFP on average after 2 hours of reaction. Compared with other reports of thermostable cell-free protein expression, our yield was comparable to 60 ng/µl obtained with the pure component CFPS system from *T. thermophilus*, while being much simpler to prepare. However, the yield was still 40% of the yield described for the reconstituted *E. coli* system (160 ng/ µl) (54), which may be caused by the degradation of mRNA by nucleases present in *T. thermophilus* cell extracts as well a temperature limit of 50 °C imposed by transcription that results in a suboptimal translation. Other thermostable CFPS based on lysates, e.g. from *T*. *kodakaraensis,* have been reported to yield 100 ng/µl after heavy optimization of the lysate preparation process and the strain used (106), unlike the solution herein proposed that relies on a wild-type strain.

Our efforts to improve the yield of the thermostable IVTT extracts included increasing the amount of RNA polymerase and *in situ* amplification of the template DNA. However, neither strategy proved fruitful (Supplementary Figures 1 and 2). Further and future endeavors to increase the yield of synthesized protein at high temperatures could include the addition of helper molecules such as chaperones, which have been shown to be beneficial and prevent aggregation, improving protein folding, solubility and functionality (107,108). While at lower temperatures, the most extensively used chaperones are *E*. *coli* trigger factor, GroES/EL and DnaK/DnaJ/GrpE chaperone systems (109,110), in *T. thermophilus* the DnaK/ClpB system and homologs of the chaperonin GroEL system can be found (111,112). Moreover, the use of chaperones, compatible solutes (113–115) or stabilizing additives such as trehalose may also have a positive effect on the insufficient stability of the thermostable T7 RNA polymerase and thus contribute to increasing the reaction temperature.

In fact, the thermostable cell-free protein synthesis failed in experiments performed above 50 °C (Figure 2), even though the optimal growth temperature of *T. thermophilus* is ca. 72 °C. The reconstituted system with purified proteins from *T. thermophilus* is functional from 37 °C to 60 °C, indicating that the translation machinery of *T. thermophilus* is active even at temperatures below the minimal growth temperature of 47 °C (55). This result agrees with our findings of slow but detectable protein synthesis at 37 °C using cell-based extracts. We attributed the upper temperature limit of our cell-free system to the tT7RNApol, which shows an optimum reaction temperature of approx. 50 °C with a half-life of 85 min at that temperature and no activity above 50-52 °C, according to the manufacturer. This upper limit could be surpassed by engineering more thermostable variants of the T7 RNA polymerase but the engineered variants reported so far cannot withstand temperatures at or above 50 °C for sustained periods of time (116). Another solution would involve the use of RNA polymerases from thermophiles or thermophilic viruses (117,118), however at the expense of the compatibility with the strong and widely used T7 promoter. For instance, the RNA polymerase the thermophilic *Geobacillus sp.* GHH01 has been described to be stable and active at elevated temperatures up to 55 °C, recognizing DNA template sequences from a wide variety of organisms, including bacteria and archaea (119), which would contribute to the applicability of thermostable IVTT extracts for recognition of diverse native promoters in the activity-based screening of metagenomic libraries.

A key issue to the viability of CFPS is the process of energy generation, which represents the major cost factor and a yield-limiting component. In fact, transcription requires thousands of nucleotide triphosphates for each mRNA transcript and translation requires two ATP equivalents for each activated tRNA molecule, two GTP equivalents per peptide bond formed and 1 GTP equivalent for the initiation and termination steps (120). With these figures, some authors have estimated the cost of batch synthesis of 1 mg/ml of a 25 kDa protein to be ca. 35– 44 mM of ATP (120). For this reason, it is logical that the productivity of batch CFPS reactions is limited by the availability of ATP (121). Generally and also in our extract composition, the supply of ATP is generated from a molecule containing high-energy phosphate bonds, such as phosphoenolpyruvate-PEP, acetyl phosphate or creatine phosphate (120), which generate ATP by a substrate-level phosphorylation reaction. However, these compounds are expensive, representing more than 50% of the total cost of the reaction (122). Moreover, they are degraded by nonspecific phosphatases present in the cell extract (120) and generate the accumulation of inorganic phosphate, which has an inhibitory effect on protein synthesis (85). These limitations can be circumvented by operating in picoliter volumes, e.g. in microfluidic droplets, which reduces the cost of reagents by orders of magnitude (61) and by using pyruvate as energy source, which produces similar protein yields as PEP at a lower cost (86), provided the multistep enzymatic reactions associated with glycolysis and oxidative phosphorylation are present (85,88,121–124). Alternatively, the use of maltodextrin in cell-free expression has been proposed in combination with polyphosphate molecules acting as phosphate donors (125), resulting in cost-effective high protein yields (71). In conclusion, developing a stable and affordable alternative supply of energy will be a key factor in increasing production rate and overall yield of the thermostable system, particularly considering the limited stability of ATP and phosphoenolpyruvate at high temperatures (126,127) and will be a determinant of the economic viability of scaled-up reactions.

For library screening applications, IVTT should ideally be driven by a single DNA molecule and produce sufficient yield to detect the final protein product. Moreover, high readout values are needed due to the fast signal integration times needed for ultrahigh-throughput operation (128). For these reasons, DNA amplification is often needed to increase the amount of template DNA within the droplets prior to the enzyme assay. Furthermore, coupling a DNA amplification step with IVTT facilitates recovery of the input DNA from a relatively low number of selected droplets after performing the screening. The isothermal conditions of RCA make it a better candidate than PCR for coupling to IVTT but it has been described that NTPs and tRNAs have an inhibitory effect on DNA replication by Φ29 DNA polymerase. Attempts to adjust the concentrations of these components and achieve a synchronous, one-pot RCA and IVTT reaction have resulted in conditions optimal for replication but still suboptimal for translation, which is critical for the sensitivity of the enzyme screening in droplets. Hence, it becomes necessary to perform the reactions sequentially: first, amplification and then, transcription-translation. Despite having attempted such a sequential setup, in this work we observed an incompatibility between the RCA reaction and *T. thermophilus*-based extracts. Our cell-free system does not work efficiently when we couple replication and transcription-translation reactions, although coupling was possible with *E. coli* extracts (Supplementary Figure 3A) or pure component IVTT mixes (65). Whether the incompatibility resides in the highly branched nature of the products (129) or in an unsuitable composition of the reaction medium is unclear. Alternative isothermal amplification methods could be considered, such as helicase-dependent Amplification (HDA)(130), recombinase polymerase amplification (RPA) (131) or the more processive T4 replisome (132).

Finally, we illustrated the applicability of *T. thermophilus* extract-based IVTT for activity-based screening by coupling cell-free expression at high temperatures with fluorescence-based activity assays. Although we have demonstrated that such a combination is possible with two different examples, background hydrolysis of substrates at higher assay temperatures will ultimately determine which enzyme activities can be successfully coupled with thermostable IVTT. As described above for the reduction of interfering nucleases, high levels of background activities can be overcome by deletion of the relevant interfering genes (133), particularly as more genome editing tools become available for thermophilic microorganisms (134). In conclusion, we have developed a cell-free expression system apt to synthesize proteins requiring high temperature for their correct folding, such as those from thermophilic organisms (135,136). Furthermore, the optimized CFPS reaction mix was amenable to coupling an enzymatic assay and is also compatible with droplet microfluidics, thus opening up the possibilities to affordably perform *in vitro* protein evolution in a cell-free context at higher temperatures, for instance to screen for thermostable protein variants for biocatalysis (137) or to perform activity-based screenings of metagenomic libraries from thermal environments. Finally, this concept can be extended to other types of extreme environments, simply by using CFPS extracts from other extremophile organisms, opening new avenues for enzyme discovery and evolution as well as synthetic biology of mesophiles and extremophiles.

## CONCLUSIONS

Currently, there are no commercial solutions to thermostable CFPS, causing researchers to make their own pure-component or cell-based extracts for IVTT. The cell-free extracts from *Thermus thermophilus* described herein represent a simpler alternative to heavily optimized or pure component thermostable *in vitro* expression systems. The protocol was simple and adaptable to different strains, with yields comparable to those of the abovementioned alternatives. Moreover, due to their compatibility with droplet microfluidics and enzyme assays at high temperatures, the reported system represents a convenient gateway for enzyme screening at higher temperatures with ultrahigh-throughput.

## DECLARATIONS

### Ethics approval and consent to participate

Not applicable

### Consent for publication

Not applicable

### Availability of Data and Materials

The datasets used and/or analysed during the current study are available from the corresponding author upon reasonable request.

### Authors’ contributions

ALJLR and PPA contributed to experimental work and were major contributors in writing the manuscript; LP and JL produced, purified and characterized the enzymatic cascade for energy regeneration; LVV, FG and MSC contributed to encapsulation of the extracts in microfluidic droplets; MSC contributed to enzymatic assays with the *T. thermophilus* BL03 S30 extracts; SC, FH, JEGP, JB and AH were responsible for securing and managing funding; FH, JEGP, JB and AH conceived the project, were responsible for writing and revising the manuscript. All authors read and approved the final manuscript.

## BIBLIOGRAPHY

1. Dondapati SK, Stech M, Zemella A, Kubick S. Cell-Free Protein Synthesis: A Promising Option for Future Drug Development. BioDrugs. 1 de junio de 2020;34(3):327-48.

2. Liu D, Evans T, Zhang F. Applications and advances of metabolite biosensors for metabolic engineering. Metabolic Engineering. septiembre de 2015;31:35–43.

3. Rogers JK, Taylor ND, Church GM. Biosensor-based engineering of biosynthetic pathways. Current Opinion in Biotechnology. diciembre de 2016;42:84–91.

4. Jiang L, Zhao J, Lian J, Xu Z. Cell-free protein synthesis enabled rapid prototyping for metabolic engineering and synthetic biology. Synthetic and Systems Biotechnology. junio de 2018;3(2):90–6.

5. Cui Z, Mureev S, Polinkovsky ME, Tnimov Z, Guo Z, Durek T, et al. Combining Sense and Nonsense Codon Reassignment for Site-Selective Protein Modification with Unnatural Amino Acids. ACS Synth Biol. 17 de marzo de 2017;6(3):535-44.

6. Cornish VW, Benson DR, Altenbach CA, Hideg K, Hubbell WL, Schultz PG. Site-specific incorporation of biophysical probes into proteins. Proc Natl Acad Sci USA. 12 de abril de 1994;91(8):2910-4.

7. Alfonta L, Zhang Z, Uryu S, Loo JA, Schultz PG. Site-Specific Incorporation of a Redox-Active Amino Acid into Proteins. J Am Chem Soc. 1 de diciembre de 2003;125(48):14662-3.

8. Neumann H, Hancock SM, Buning R, Routh A, Chapman L, Somers J, et al. A Method for Genetically Installing Site-Specific Acetylation in Recombinant Histones Defines the Effects of H3 K56 Acetylation. Molecular Cell. octubre de 2009;36(1):153–63.

9. Nguyen DP, Garcia Alai MM, Kapadnis PB, Neumann H, Chin JW. Genetically Encoding N ^ɛ^ -Methyl-L-lysine in Recombinant Histones. J Am Chem Soc. 14 de octubre de 2009;131(40):14194-5.

10. Virdee S, Kapadnis PB, Elliott T, Lang K, Madrzak J, Nguyen DP, et al. Traceless and Site-Specific Ubiquitination of Recombinant Proteins. J Am Chem Soc. 20 de julio de 2011;133(28):10708-11.

11. Oza JP, Aerni HR, Pirman NL, Barber KW, ter Haar CM, Rogulina S, et al. Robust production of recombinant phosphoproteins using cell-free protein synthesis. Nat Commun. 9 de septiembre de 2015;6(1):8168.

12. Boyer ME, Wang CW, Swartz JR. Simultaneous expression and maturation of the iron-sulfur protein ferredoxin in a cell-free system. Biotechnol Bioeng. 5 de mayo de 2006;94(1):128-38.

13. Kuchenreuther JM, Shiigi SA, Swartz JR. Cell-Free Synthesis of the H-Cluster: A Model for the In Vitro Assembly of Metalloprotein Metal Centers. En: Fontecilla-Camps JC, Nicolet Y, editores. Metalloproteins [Internet]. Totowa, NJ: Humana Press; 2014 [citado 22 de marzo de 2023]. p. 49-72. (Methods in Molecular Biology; vol. 1122). Disponible en: http://link.springer.com/10.1007/978-1-62703-794-5_5

14. Kigawa T, Yamaguchi-Nunokawa E, Kodama K, Matsuda T, Yabuki T, Matsuda N, et al. [No title found]. Journal of Structural and Functional Genomics. 2002;2(1):29–35.

15. Karim AS, Dudley QM, Juminaga A, Yuan Y, Crowe SA, Heggestad JT, et al. In vitro prototyping and rapid optimization of biosynthetic enzymes for cell design. Nat Chem Biol. agosto de 2020;16(8):912–9.

16. Kuruma Y, Stano P, Ueda T, Luisi PL. A synthetic biology approach to the construction of membrane proteins in semi-synthetic minimal cells. Biochimica et Biophysica Acta (BBA) - Biomembranes. febrero de 2009;1788(2):567–74.

17. Wu F, Tan C. The engineering of artificial cellular nanosystems using synthetic biology approaches: Artificial cellular nanosystems using synthetic biology approaches. WIREs Nanomed Nanobiotechnol. julio de 2014;6(4):369–83.

18. Elani Y, Law RV, Ces O. Protein synthesis in artificial cells: using compartmentalisation for spatial organisation in vesicle bioreactors. Phys Chem Chem Phys. 2015;17(24):15534–7.

19. He M, Taussig MJ. DiscernArray^TM^ technology: a cell-free method for the generation of protein arrays from PCR DNA. Journal of Immunological Methods. marzo de 2003;274(1-2):265–70.

20. Angenendt P, Nyarsik L, Szaflarski W, Glökler J, Nierhaus KH, Lehrach H, et al. Cell-Free Protein Expression and Functional Assay in Nanowell Chip Format. Anal Chem. 1 de abril de 2004;76(7):1844-9.

21. Pardee K, Green AA, Ferrante T, Cameron DE, DaleyKeyser A, Yin P, et al. Paper-Based Synthetic Gene Networks. Cell. noviembre de 2014;159(4):940–54.

22. Kelwick RJR, Webb AJ, Wang Y, Heliot A, Allan F, Emery AM, et al. AL-PHA beads: Bioplastic-based protease biosensors for global health applications. Materials Today. julio de 2021;47:25–37.

23. Hicks M, Bachmann TT, Wang B. Synthetic Biology Enables Programmable Cell-Based Biosensors. ChemPhysChem. 16 de enero de 2020;21(2):132-44.

24. Karim AS, Jewett MC. A cell-free framework for rapid biosynthetic pathway prototyping and enzyme discovery. Metabolic Engineering. julio de 2016;36:116–26.

25. Wu YY, Culler S, Khandurina J, Van Dien S, Murray RM. Prototyping 1,4-butanediol (BDO) biosynthesis pathway in a cell-free transcription-translation (TX-TL) system [Internet]. Synthetic Biology; 2015 abr [citado 12 de mayo de 2023]. Disponible en: http://biorxiv.org/lookup/doi/10.1101/017814

26. Wu YY, Sato H, Huang H, Culler SJ, Khandurina J, Nagarajan H, et al. System-level studies of a cell-free transcription-translation platform for metabolic engineering [Internet]. Bioengineering; 2017 ago [citado 12 de mayo de 2023]. Disponible en: http://biorxiv.org/lookup/doi/10.1101/172007

27. Savage DF, Anderson CL, Robles-Colmenares Y, Newby ZE, Stroud RM. Cell-free complements in vivo expression of the E. coli membrane proteome. Protein Sci. mayo de 2007;16(5):966–76.

28. Schwarz D, Daley D, Beckhaus T, Dötsch V, Bernhard F. Cell-free expression profiling of E. coli inner membrane proteins. Proteomics. mayo de 2010;10(9):1762–79.

29. Yang JP, Cirico T, Katzen F, Peterson TC, Kudlicki W. Cell-free synthesis of a functional G protein-coupled receptor complexed with nanometer scale bilayer discs. BMC Biotechnol. diciembre de 2011;11(1):57.

30. Martemyanov KA, Shirokov VA, Kurnasov OV, Gudkov AT, Spirin AS. Cell-Free Production of Biologically Active Polypeptides: Application to the Synthesis of Antibacterial Peptide Cecropin. Protein Expression and Purification. abril de 2001;21(3):456–61.

31. Zawada JF, Yin G, Steiner AR, Yang J, Naresh A, Roy SM, et al. Microscale to manufacturing scale-up of cell-free cytokine production—a new approach for shortening protein production development timelines. Biotechnol Bioeng. julio de 2011;108(7):1570–8.

32. Lu Y, Welsh JP, Swartz JR. Production and stabilization of the trimeric influenza hemagglutinin stem domain for potentially broadly protective influenza vaccines. Proc Natl Acad Sci USA. 7 de enero de 2014;111(1):125-30.

33. Yin G, Garces ED, Yang J, Zhang J, Tran C, Steiner AR, et al. Aglycosylated antibodies and antibody fragments produced in a scalable in vitro transcription-translation system. mAbs. marzo de 2012;4(2):217–25.

34. Spirin AS, Baranov VI, Ryabova LA, Ovodov S, Alakhov YB. A Continuous Cell-Free Translation System Capable of Producing Polypeptides in High Yield. Science. 25 de noviembre de 1988;242(4882):1162-4.

35. Endo Y, Sawasaki T. High-throughput, genome-scale protein production method based on the wheat germ cell-free expression system. Biotechnology Advances. noviembre de 2003;21(8):695–713.

36. Harbers M. Wheat germ systems for cell-free protein expression. FEBS Letters. 25 de agosto de 2014;588(17):2762-73.

37. Sissons CH. Yeast protein synthesis. Preparation and analysis of a highly active cell-free system. Biochemical Journal. 1 de octubre de 1974;144(1):131-40.

38. Gallis BM, McDonnell JP, Hopper JE, Young ET. Translation of poly(riboadenylic acid)-enriched messenger RNAs from the yeast, Saccharomyces cerevisiae, in heterologous cell-free systems. Biochemistry. 11 de marzo de 1975;14(5):1038-46.

39. Hempel R, Schmidt-Brauns J, Gebinoga M, Wirsching F, Schwienhorst A. Cation Radius Effects on Cell-Free Translation in Rabbit Reticulocyte Lysate. Biochemical and Biophysical Research Communications. mayo de 2001;283(2):267–72.

40. Weber LA, Feman ER, Baglioni C. Cell free system from HeLa cells active in initiation of protein synthesis. Biochemistry. 1 de diciembre de 1975;14(24):5315-21.

41. Moore SJ, Lai HE, Needham H, Polizzi KM, Freemont PS. Streptomyces venezuelae TX-TL - a next generation cell-free synthetic biology tool. Biotechnology Journal. abril de 2017;12(4):1600678.

42. Li J, Wang H, Kwon YC, Jewett MC. Establishing a high yielding streptomyces -based cell-free protein synthesis system: Establishing a Streptomyces -Based CFPS System. Biotechnol Bioeng. junio de 2017;114(6):1343–53.

43. Kelwick R, Webb AJ, MacDonald JT, Freemont PS. Development of a Bacillus subtilis cell-free transcription-translation system for prototyping regulatory elements. Metabolic Engineering. noviembre de 2016;38:370–81.

44. Des Soye BJ, Davidson SR, Weinstock MT, Gibson DG, Jewett MC. Establishing a High-Yielding Cell-Free Protein Synthesis Platform Derived from Vibrio natriegens. ACS Synth Biol. 21 de septiembre de 2018;7(9):2245-55.

45. Failmezger J, Scholz S, Blombach B, Siemann-Herzberg M. Cell-Free Protein Synthesis From Fast-Growing Vibrio natriegens. Front Microbiol. 1 de junio de 2018;9:1146.

46. Wiegand DJ, Lee HH, Ostrov N, Church GM. Establishing a cell-free Vibrio natriegens expression system [Internet]. Synthetic Biology; 2018 may [citado 22 de marzo de 2023]. Disponible en: http://biorxiv.org/lookup/doi/10.1101/331645

47. Dondapati SK, Kreir M, Quast RB, Wüstenhagen DA, Brüggemann A, Fertig N, et al. Membrane assembly of the functional KcsA potassium channel in a vesicle-based eukaryotic cell-free translation system. Biosensors and Bioelectronics. septiembre de 2014;59:174–83.

48. Quast RB, Sonnabend A, Stech M, Wüstenhagen DA, Kubick S. High-yield cell-free synthesis of human EGFR by IRES-mediated protein translation in a continuous exchange cell-free reaction format. Sci Rep. 26 de julio de 2016;6(1):30399.

49. Mikami S, Masutani M, Sonenberg N, Yokoyama S, Imataka H. An efficient mammalian cell-free translation system supplemented with translation factors. Protein Expression and Purification. abril de 2006;46(2):348–57.

50. Buntru M, Vogel S, Stoff K, Spiegel H, Schillberg S. A versatile coupled cell-free transcription-translation system based on tobacco BY-2 cell lysates: A Coupled Cell-free Transcription-Translation System. Biotechnol Bioeng. mayo de 2015;112(5):867–78.

51. Kovtun O, Mureev S, Johnston W, Alexandrov K. Towards the Construction of Expressed Proteomes Using a Leishmania tarentolae Based Cell-Free Expression System. Preiss T, editor. PLoS ONE. 21 de diciembre de 2010;5(12):e14388.

52. Kovtun O, Mureev S, Jung W, Kubala MH, Johnston W, Alexandrov K. Leishmania cell-free protein expression system. Methods. septiembre de 2011;55(1):58–64.

53. Yim SS, Johns NI, Park J, Gomes AL, McBee RM, Richardson M, et al. Multiplex transcriptional characterizations across diverse bacterial species using cell-free systems. Mol Syst Biol [Internet]. agosto de 2019 [citado 29 de marzo de 2023];15(8). Disponible en: https://onlinelibrary.wiley.com/doi/10.15252/msb.20198875

54. Shimizu Y, Inoue A, Tomari Y, Suzuki T, Yokogawa T, Nishikawa K, et al. Cell-free translation reconstituted with purified components. Nat Biotechnol. agosto de 2001;19(8):751–5.

55. Zhou Y, Asahara H, Gaucher EA, Chong S. Reconstitution of translation from Thermus thermophilus reveals a minimal set of components sufficient for protein synthesis at high temperatures and functional conservation of modern and ancient translation components. Nucleic Acids Research. 1 de septiembre de 2012;40(16):7932-45.

56. Cai Q, Hanson JA, Steiner AR, Tran C, Masikat MR, Chen R, et al. A simplified and robust protocol for immunoglobulin expression in E scherichia coli cell-free protein synthesis systems. Biotechnol Progress. mayo de 2015;31(3):823–31.

57. Garamella J, Marshall R, Rustad M, Noireaux V. The All E. coli TX-TL Toolbox 2.0: A Platform for Cell-Free Synthetic Biology. ACS Synth Biol. 15 de abril de 2016;5(4):344-55.

58. Myers TW, Gelfand DH. Reverse transcription and DNA amplification by a Thermus thermophilus DNA polymerase. Biochemistry. 1 de agosto de 1991;30(31):7661-6.

59. Ruggero D, Creti R, Londei P. In vitro translation of archaeal natural mRNAs at high temperature. FEMS Microbiology Letters. febrero de 1993;107(1):89–94.

60. Ohno-Iwashita Y, Oshima T, Imahori K. In vitro protein synthesis at elevated temperature by an extract of an extreme thermophile: Effects of polyamines on the polyuridylic acid-directed reaction. Archives of Biochemistry and Biophysics. 1975;171(2):490–9.

61. Agresti JJ, Antipov E, Abate AR, Ahn K, Rowat AC, Baret JC, et al. Ultrahigh-throughput screening in drop-based microfluidics for directed evolution. Proc Natl Acad Sci USA. 2 de marzo de 2010;107(9):4004-9.

62. Tawfik DS, Griffiths AD. Man-made cell-like compartments for molecular evolution. Nat Biotechnol. julio de 1998;16(7):652–6.

63. Griffiths AD. Directed evolution of an extremely fast phosphotriesterase by in vitro compartmentalization. The EMBO Journal. 2 de enero de 2003;22(1):24-35.

64. Aharoni A, Thieme K, Chiu CPC, Buchini S, Lairson LL, Chen H, et al. High-throughput screening methodology for the directed evolution of glycosyltransferases. Nat Methods. agosto de 2006;3(8):609–14.

65. Holstein JM, Gylstorff C, Hollfelder F. Cell-free Directed Evolution of a Protease in Microdroplets at Ultrahigh Throughput. ACS Synth Biol. 19 de febrero de 2021;10(2):252-7.

66. Schaerli Y, Wootton RC, Robinson T, Stein V, Dunsby C, Neil MAA, et al. Continuous-Flow Polymerase Chain Reaction of Single-Copy DNA in Microfluidic Microdroplets. Anal Chem. 1 de enero de 2009;81(1):302-6.

67. Diamante L, Gatti-Lafranconi P, Schaerli Y, Hollfelder F. In vitro affinity screening of protein and peptide binders by megavalent bead surface display. Protein Engineering Design and Selection. 1 de octubre de 2013;26(10):713-24.

68. Leis B, Angelov A, Li H, Liebl W. Genetic analysis of lipolytic activities in Thermus thermophilus HB27. Journal of Biotechnology. 10 de diciembre de 2014;191:150-7.

69. Hanahan D. Studies on transformation of Escherichia coli with plasmids. Journal of Molecular Biology. junio de 1983;166(4):557–80.

70. Sun ZZ, Hayes CA, Shin J, Caschera F, Murray RM, Noireaux V. Protocols for Implementing an Escherichia coli Based TX-TL Cell-Free Expression System for Synthetic Biology. JoVE. 16 de septiembre de 2013;(79):50762.

71. Caschera F, Noireaux V. Preparation of amino acid mixtures for cell-free expression systems. BioTechniques. enero de 2015;58(1):40–3.

72. Laemmli UK. Cleavage of Structural Proteins during the Assembly of the Head of Bacteriophage T4. Nature. agosto de 1970;227(5259):680-5.

73. Schindelin J, Arganda-Carreras I, Frise E, Kaynig V, Longair M, Pietzsch T, et al. Fiji: an open-source platform for biological-image analysis. Nat Methods. julio de 2012;9(7):676-82.

74. Coquelle N, Fioravanti E, Weik M, Vellieux F, Madern D. Activity, Stability and Structural Studies of Lactate Dehydrogenases Adapted to Extreme Thermal Environments. Journal of Molecular Biology. noviembre de 2007;374(2):547–62.

75. Arai A, Murakami S, Nakajima M o. Purification and Characterization of a Thermostable Pyruvate Kinase from the Actinomycete Microbispora thermodiastatica. Bioscience, Biotechnology, and Biochemistry. enero de 1997;61(1):40–5.

76. Rusnak P, Haney P, Konisky J. The adenylate kinases from a mesophilic and three thermophilic methanogenic members of the Archaea. J Bacteriol. junio de 1995;177(11):2977–81.

77. Verhoeven JA, Schenck KM, Meyer RR, Trela JM. Purification and characterization of an inorganic pyrophosphatase from the extreme thermophile Thermus aquaticus. J Bacteriol. octubre de 1986;168(1):318–21.

78. Chan KM, Delfert D, Junger KD. A direct colorimetric assay for Ca2+-stimulated ATPase activity. Analytical Biochemistry. septiembre de 1986;157(2):375–80.

79. Fernández-Herrero L A, Olabarría G, Castón J R, Lasa I, Berenguer J. Horizontal transference of S-layer genes within Thermus thermophilus. Journal of Bacteriology. 1 de octubre de 1995;177(19):5460-6.

80. Kim TW, Kim DM, Choi CY. Rapid production of milligram quantities of proteins in a batch cell-free protein synthesis system. Journal of Biotechnology. julio de 2006;124(2):373–80.

81. Liu DV, Zawada JF, Swartz JR. Streamlining Escherichia Coli S30 Extract Preparation for Economical Cell-Free Protein Synthesis. Biotechnol Progress. 5 de septiembre de 2008;21(2):460-5.

82. Gąciarz A, Khatri NK, Velez-Suberbie ML, Saaranen MJ, Uchida Y, Keshavarz-Moore E, et al. Efficient soluble expression of disulfide bonded proteins in the cytoplasm of Escherichia coli in fed-batch fermentations on chemically defined minimal media. Microb Cell Fact. diciembre de 2017;16(1):108.

83. Jackson AM. Cell-free protein synthesis for proteomics. Briefings in Functional Genomics and Proteomics. 1 de enero de 2004;2(4):308-19.

84. Uzawa T, Yamagishi A, Oshima T. Continuous Cell-free Protein Synthesis Directed by Messenger DNA and Catalyzed by Extract of Thermus thermophilus HB27. Bioscience, Biotechnology, and Biochemistry. enero de 2003;67(3):639–42.

85. Kim DM, Swartz JR. Regeneration of adenosine triphosphate from glycolytic intermediates for cell-free protein synthesis. Biotechnol Bioeng. 20 de agosto de 2001;74(4):309-16.

86. Jewett MC, Swartz JR. Mimicking theEscherichia coli cytoplasmic environment activates long-lived and efficient cell-free protein synthesis. Biotechnol Bioeng. 5 de abril de 2004;86(1):19-26.

87. Kim DM, Kigawa T, Choi CY, Yokoyama S. A Highly Efficient Cell-Free Protein Synthesis System from Escherichia coli. Eur J Biochem. agosto de 1996;239(3):881–6.

88. Sitaraman K, Esposito D, Klarmann G, Le Grice SF, Hartley JL, Chatterjee DK. A novel cell-free protein synthesis system. Journal of Biotechnology. junio de 2004;110(3):257–63.

89. Shin J, Noireaux V. Efficient cell-free expression with the endogenous E. Coli RNA polymerase and sigma factor 70. J Biol Eng. 2010;4(1):8.

90. Cava F, de Pedro MA, Blas-Galindo E, Waldo GS, Westblade LF, Berenguer J. Expression and use of superfolder green fluorescent protein at high temperatures in vivo: a tool to study extreme thermophile biology. Environ Microbiol. marzo de 2008;10(3):605–13.

91. Dean FB, Nelson JR, Giesler TL, Lasken RS. Rapid Amplification of Plasmid and Phage DNA Using Phi29 DNA Polymerase and Multiply-Primed Rolling Circle Amplification. Genome Res. 1 de junio de 2001;11(6):1095-9.

92. Sandoval M, Ferreras E, Pérez-Sánchez M, Berenguer J, Sinisterra JV, Hernaiz MJ. Screening of strains and recombinant enzymes from Thermus thermophilus for their use in disaccharide synthesis. Journal of Molecular Catalysis B: Enzymatic. febrero de 2012;74(3-4):162–9.

93. Vieille C, Zeikus GJ. Hyperthermophilic Enzymes: Sources, Uses, and Molecular Mechanisms for Thermostability. Microbiol Mol Biol Rev. marzo de 2001;65(1):1–43.

94. Yang HL, Ivashkiv L, Chen HZ, Zubay G, Cashel M. Cell-free coupled transcription-translation system for investigation of linear DNA segments. Proc Natl Acad Sci USA. diciembre de 1980;77(12):7029–33.

95. Bassett CL, Rawson JR. In vitro coupled transcription-translation of linear DNA fragments in a lysate derived from a recB rna pnp strain of Escherichia coli. J Bacteriol. diciembre de 1983;156(3):1359–62.

96. Dillingham MS, Kowalczykowski SC. RecBCD Enzyme and the Repair of Double-Stranded DNA Breaks. Microbiol Mol Biol Rev. diciembre de 2008;72(4):642–71.

97. Lehman IR. The deoxyribonucleases of Escherichia coli. I. Purification and properties of a phosphodiesterase. J Biol Chem. mayo de 1960;235:1479–87.

98. Batista AC, Levrier A, Soudier P, Voyvodic PL, Achmedov T, Reif-Trauttmansdorff T, et al. Differentially Optimized Cell-Free Buffer Enables Robust Expression from Unprotected Linear DNA in Exonuclease-Deficient Extracts. ACS Synth Biol. 18 de febrero de 2022;11(2):732-46.

99. Sabeti Azad M, Cardoso Batista A, Faulon JL, Beisel CL, Bonnet J, Kushwaha M. Cell-Free Protein Synthesis from Exonuclease-Deficient Cellular Extracts Utilizing Linear DNA Templates. JoVE. 9 de agosto de 2022;(186):64236.

100. Verdú C, Pérez-Arnaiz P, Peropadre A, Berenguer J, Mencía M. Deletion of the primase-polymerases encoding gene, located in a mobile element in Thermus thermophilus HB27, leads to loss of function mutation of addAB genes. Frontiers in Microbiology [Internet]. 2022;13. Disponible en: https://www.frontiersin.org/articles/10.3389/fmicb.2022.1005862

101. Arce A, Guzman Chavez F, Gandini C, Puig J, Matute T, Haseloff J, et al. Decentralizing Cell-Free RNA Sensing With the Use of Low-Cost Cell Extracts. Front Bioeng Biotechnol. 23 de agosto de 2021;9:727584.

102. Sun ZZ, Yeung E, Hayes CA, Noireaux V, Murray RM. Linear DNA for Rapid Prototyping of Synthetic Biological Circuits in an Escherichia coli Based TX-TL Cell-Free System. ACS Synth Biol. 20 de junio de 2014;3(6):387-97.

103. Seki E, Matsuda N, Yokoyama S, Kigawa T. Cell-free protein synthesis system from Escherichia coli cells cultured at decreased temperatures improves productivity by decreasing DNA template degradation. Analytical Biochemistry. junio de 2008;377(2):156–61.

104. Yim SS, Johns NI, Noireaux V, Wang HH. Protecting Linear DNA Templates in Cell-Free Expression Systems from Diverse Bacteria. ACS Synth Biol. 16 de octubre de 2020;9(10):2851-5.

105. Norouzi M, Panfilov S, Pardee K. High-Efficiency Protection of Linear DNA in Cell-Free Extracts from Escherichia coli and Vibrio natriegens. ACS Synth Biol. 16 de julio de 2021;10(7):1615-24.

106. Endoh T, Kanai T, Imanaka T. A highly productive system for cell-free protein synthesis using a lysate of the hyperthermophilic archaeon, Thermococcus kodakaraensis. Appl Microbiol Biotechnol. abril de 2007;74(5):1153–61.

107. Ryabova LA, Desplancq D, Spirin AS, Plückthun A. Functional antibody production using cell-free translation: Effects of protein disulfide isomerase and chaperones. Nat Biotechnol. enero de 1997;15(1):79–84.

108. Jiang X, Ookubo Y, Fujii I, Nakano H, Yamane T. Expression of Fab fragment of catalytic antibody 6D9 in an Escherichia coli in vitro coupled transcription/translation system. FEBS Letters. 13 de marzo de 2002;514(2-3):290-4.

109. Fink AL. Chaperone-Mediated Protein Folding. Physiological Reviews. 1 de abril de 1999;79(2):425-49.

110. Niwa T, Kanamori T, Ueda T, Taguchi H. Global analysis of chaperone effects using a reconstituted cell-free translation system. Proc Natl Acad Sci USA. 5 de junio de 2012;109(23):8937-42.

111. A chaperonin from a thermophilic bacterium, Thermus thermophilus. Phil Trans R Soc Lond B. 29 de marzo de 1993;339(1289):305-12.

112. Schlee S, Reinstein J. The DnaK/ClpB chaperone system from Thermus thermophilus. CMLS, Cell Mol Life Sci. octubre de 2002;59(10):1598–606.

113. Borges N, Ramos A, Raven N, Sharp R, Santos H. Comparative study of the thermostabilizing properties of mannosylglycerate and other compatible solutes on model enzymes. Extremophiles. 1 de junio de 2002;6(3):209-16.

114. Carninci P, Nishiyama Y, Westover A, Itoh M, Nagaoka S, Sasaki N, et al. Thermostabilization and thermoactivation of thermolabile enzymes by trehalose and its application for the synthesis of full length cDNA. Proc Natl Acad Sci USA. 20 de enero de 1998;95(2):520-4.

115. Ramos A, Raven N, Sharp RJ, Bartolucci S, Rossi M, Cannio R, et al. Stabilization of Enzymes against Thermal Stress and Freeze-Drying by Mannosylglycerate. Appl Environ Microbiol. octubre de 1997;63(10):4020–5.

116. Wu MZ, Asahara H, Tzertzinis G, Roy B. Synthesis of low immunogenicity RNA with high-temperature in vitro transcription. RNA. marzo de 2020;26(3):345–60.

117. Doss RK, Palmer M, Mead DA, Hedlund BP. Functional biology and biotechnology of thermophilic viruses. Essays in Biochemistry. 11 de agosto de 2023;67(4):671-84.

118. Yu MX, Slater MR, Ackermann HW. Isolation and characterization of Thermus bacteriophages. Archives of Virology. 1 de abril de 2006;151(4):663-79.

119. Kinfu BM, Jahnke M, Janus M, Besirlioglu V, Roggenbuck M, Meurer R, et al. Recombinant RNA Polymerase from Geobacillus sp. GHH01 as tool for rapid generation of metagenomic RNAs using in vitro technologies. Biotechnology and Bioengineering. diciembre de 2017;114(12):2739-52.

120. Calhoun KA, Swartz JR. Energy Systems for ATP Regeneration in Cell-Free Protein Synthesis Reactions. En: Grandi G, editor. In Vitro Transcription and Translation Protocols [Internet]. Totowa, NJ: Humana Press; 2007 [citado 28 de marzo de 2023]. p. 3-17. Disponible en: http://link.springer.com/10.1007/978-1-59745-388-2_1

121. Kim DM, Swartz JR. Prolonging cell-free protein synthesis with a novel ATP regeneration system. Biotechnol Bioeng. 1999;66(3):180–8.

122. Kim TW, Oh IS, Keum JW, Kwon YC, Byun JY, Lee KH, et al. Prolonged cell-free protein synthesis using dual energy sources: Combined use of creatine phosphate and glucose for the efficient supply of ATP and retarded accumulation of phosphate. Biotechnol Bioeng. 15 de agosto de 2007;97(6):1510-5.

123. Kim HC, Kim DM. Methods for energizing cell-free protein synthesis. Journal of Bioscience and Bioengineering. julio de 2009;108(1):1–4.

124. Calhoun KA, Swartz JR. Energizing cell-free protein synthesis with glucose metabolism. Biotechnol Bioeng. 5 de junio de 2005;90(5):606-13.

125. Wang Y, Zhang YHP. Cell-free protein synthesis energized by slowly-metabolized maltodextrin. BMC Biotechnol. 2009;9(1):58.

126. Kengen ServéWM, Stams AJM, De Vos WM. Sugar metabolism of hyperthermophiles. FEMS Microbiol Rev. mayo de 1996;18(2-3):119–37.

127. Schramm A, Siebers B, Tjaden B, Brinkmann H, Hensel R. Pyruvate Kinase of the Hyperthermophilic Crenarchaeote Thermoproteus tenax: Physiological Role and Phylogenetic Aspects. J Bacteriol. abril de 2000;182(7):2001–9.

128. Mair P, Gielen F, Hollfelder F. Exploring sequence space in search of functional enzymes using microfluidic droplets. Current Opinion in Chemical Biology. abril de 2017;37:137–44.

129. Zhang Y, Tanner NA. Isothermal Amplification of Long, Discrete DNA Fragments Facilitated by Single-Stranded Binding Protein. Sci Rep. 17 de agosto de 2017;7(1):8497.

130. Vincent M, Xu Y, Kong H. Helicase-dependent isothermal DNA amplification. EMBO Rep. agosto de 2004;5(8):795–800.

131. Piepenburg O, Williams CH, Stemple DL, Armes NA. DNA Detection Using Recombination Proteins. Haber J, editor. PLoS Biol. 13 de junio de 2006;4(7):e204.

132. Schaerli Y, Stein V, Spiering MM, Benkovic SJ, Abell C, Hollfelder F. Isothermal DNA amplification using the T4 replisome: circular nicking endonuclease-dependent amplification and primase-based whole-genome amplification. Nucleic Acids Research. 1 de diciembre de 2010;38(22):e201-e201.

133. Leis B, Angelov A, Li H, Liebl W. Genetic analysis of lipolytic activities in Thermus thermophilus HB27. Journal of Biotechnology. diciembre de 2014;191:150–7.

134. Adalsteinsson BT, Kristjansdottir T, Merre W, Helleux A, Dusaucy J, Tourigny M, et al. Efficient genome editing of an extreme thermophile, Thermus thermophilus, using a thermostable Cas9 variant. Sci Rep. 5 de mayo de 2021;11(1):9586.

135. Hidalgo A, Betancor L, Moreno R, Zafra O, Cava F, Fernández-Lafuente R, et al. Thermus thermophilus as a cell factory for the production of a thermophilic Mn-dependent catalase which fails to be synthesized in an active form in Escherichia coli. Appl Environ Microbiol. julio de 2004;70(7):3839–44.

136. Ribeiro AL, Sánchez M, Hidalgo A, Berenguer J. Stabilization of Enzymes by Using Thermophiles. En: Barredo JL, Herráiz I, editores. Microbial Steroids [Internet]. New York, NY: Springer New York; 2017 [citado 29 de marzo de 2023]. p. 297-312. (Methods in Molecular Biology; vol. 1645). Disponible en: http://link.springer.com/10.1007/978-1-4939-7183-1_21

137. Bommarius AS, Broering JM, Chaparro-Riggers JF, Polizzi KM. High-throughput screening for enhanced protein stability. Current Opinion in Biotechnology. diciembre de 2006;17(6):606–10.

