## Supplementary material for "Thermostable *in vitro* transcription-translation for enzyme screening in microdroplets"

Supplementary Table 1. Plasmids used in this work

| **Plasmid** | **Description** | **Reference** |
|---|---|---|
| pET28b(+) | Vector for gene expression | Novagen |
| pET28b_sGFP | Vector for sGFP expression | This work |
| pET22b_*TTP0042* | Vector for TT_P0042 expression | (92) |
| PET28b_Bst | Vector for esterase BstE expression | This work |
| PET28b _PstE | Vector for esterase PestE expression | This work |

Supplementary Table 2. Primers used in this work.

| **Primer** | **Description** | **Sequence (5' > 3')** |
|---|---|---|
| PK_NdeI_Fw | Amplification of PK | TTTTTTCATATGCCGCCTTTTAAGCG |
| PK_HindIII_Rv | Amplification of PK | TTTTTAAGCTTCCCCACCCGCTCCA |
| NDK_NdeI_Fw | Amplification of NDK | TTTTTTCATATGGAGCGGACCTTCG |
| NDK_HindIII_Rv | Amplification of NDK | TTTTTTAAGCTTAAGGAGCTCCTCGG |
| ADK_NdeI_Fw | Amplification of ADK | TTTTTTCATATGGTGGACGTGGGACA |
| ADK_EcoRI_Rv | Amplification of ADK | TTTTTTAAGCTTGATCCCTAACGCCG |
| LDH_NdeI_fw | Amplification of LDH | AAAAAACATATGAAGGTCGGCATCGTG |
| LDH_HindIII_rv | Amplification of LDH | AAAAAAGCTTCTAAAACCCCAGGGCGAAGGCCGCC |
| PPI_NdeI_Fw | Amplification of IPP | TTTTTTCATATGGCGAACCTGAAGAG |
| PPI_HindIII_Rv | Amplification of IPP | TTTTTAAGCTTGCCCTTGTAGCGGG |
| Bst_NdeI_Fw | Amplification of Bst | AAAAAACATATGATGAAAATCGTTCC |
| Bst_stop_HindIII_Rv | Amplification of Bst | AAAAAAAAGCTTTTACCAATCTAACGATTC |
| PstE_NdeI_Fw | Amplification of PstE | AAAAAACATATGCCGCTGAGCCCG |
| PstE_stop_HindIII_Rv | Amplification of PstE | AAAAAAAAGCTTTTACGCCACAGCCATC |
| T7_prom_Fw | Sequencing and amplification of sGFP from pET plasmids | TAATACGACTCACTATAGGG |

| | | |
|---|---|---|
| T7_term_Rv | Sequencing and amplification of sGFP from pET plasmids | GCTAGTTATTGCTCAGCGG |

Supplementary Table 3. Activity and melting temperature of *T. thermophilus* energy regeneration enzymes

| | **Activity (U/mg)** | **$T_m$ (°C)** |
|---|---|---|
| **PK** | 86.1 | 93.5 |
| **NDK** | 11.8 | ≥100 |
| **ADK** | 110.6 | 97.6 |
| **IPP** | 0.72 | 92.0 |

**SUPPLEMENTARY FIGURE LEGENDS**

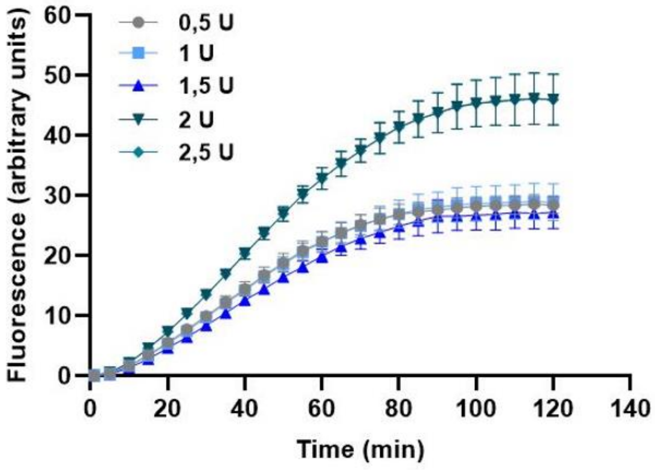


**Supplementary Figure 1. Synthesis of sGFP in the presence of different amounts of thermostable T7 RNA polymerase.** Reaction mixtures containing 40 ng/µl of pET28b_sGFP and the indicated concentration (0.5, 1, 1.5, 2, 2.5 U/µl of reaction) of thermostable T7 RNA polymerase were incubated at 50 °C for 120 min. Composition of the reaction mixtures are indicated in Table 1. The amount of protein synthesized was monitored in real time as fluorescence emission. Results are the average of n=3 reactions and error bars represent standard deviations.

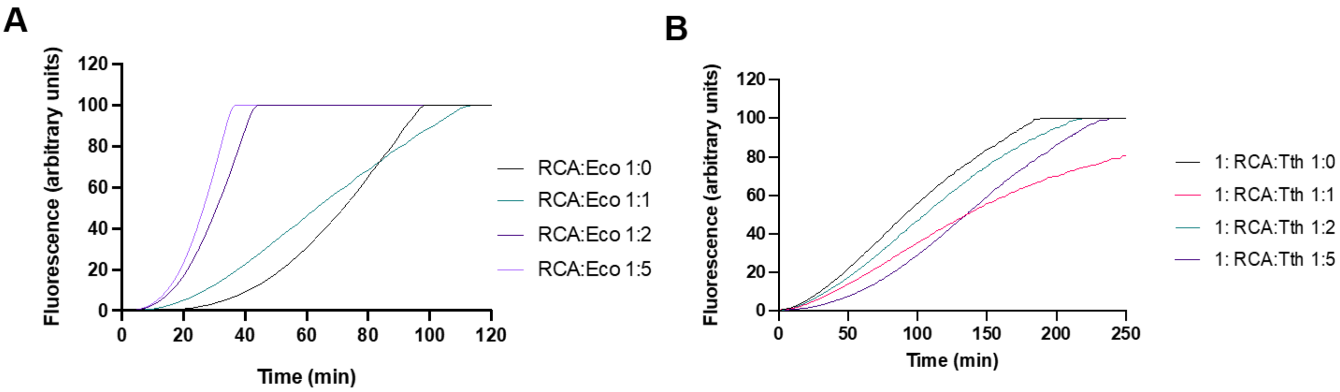


**Supplementary Figure 2. Coupling non-simultaneous, DNA amplification, transcription and translation reactions. A.** Reaction mixtures containing 1 ng of pET28b_*sGFP* were incubated at 30 °C for 180 min for RCA-DNA amplification, then IVTT components from *E. coli*

(PURExpress®, NEB) were added at different ratios and incubated at 37 °C for 180 min more. **B.** Same initial reaction mixture as in A After RCA-DNA amplification, *T. thermophilus* extracts were added at different ratios and incubated at 50 °C for 180 min. As negative control, reactions in A and B were performed in which no RCA components were added. Composition of the reaction mixtures are indicated in Table 1. sGFP synthesized was monitored in real time as fluorescence emission.

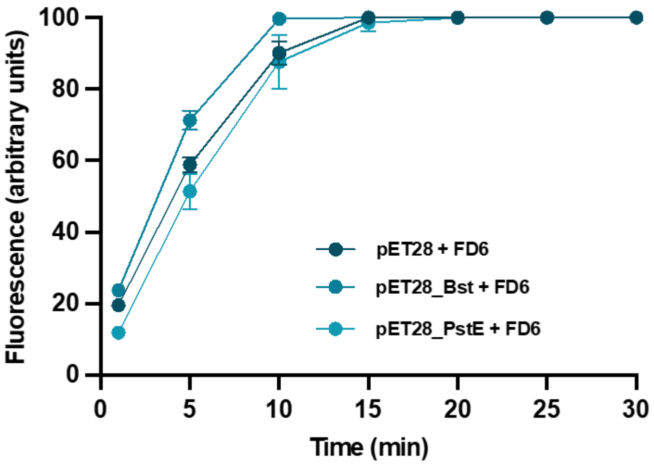


**Supplementary Figure 3. Simultaneous cell-free one-pot synthesis and esterase activity assay at different temperatures.** Reaction mixtures containing 40 ng/µl of pET28.His.TEV_Bst or pET28.His.TEV_PstE and 5 µM of fluorogenic substrate FD6 were incubated at 50 °C for 30 min. As negative controls of the experiment, an empty pET28b plasmid was incubated in the presence of FD6. Composition of the reaction mixtures are indicated in Table 3. Results are the average of n=3 reactions and error bars represent standard deviations.

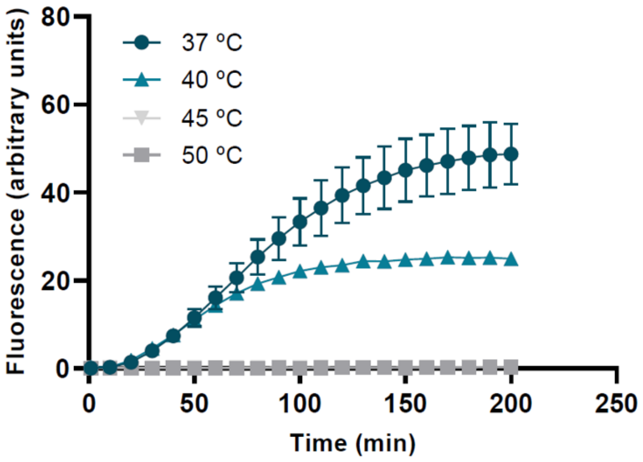

**Supplementary Figure 4. Synthesis of sGFP using PURExpress® at different temperatures.** Reaction mixtures containing 5 ng of pET28_*sGFP* as template were incubated at 37 ºC (circles), 40 ºC (triangles), 45 ºC (inverted triangles) and 50 ºC (squares) for 200 min for *in vitro* transcription and translation using reconstituted *E. coli* components (PURExpress®). sGFP synthesized was monitored in real time as fluorescence emission.
